# Insights into Heart Failure Metabolite Markers through Explainable Machine Learning

**DOI:** 10.1101/2024.10.04.616718

**Authors:** Cantin Baron, Pamela Mehanna, Caroline Daneault, Leslie Hausermann, David Busseuil, Jean-Claude Tardif, Jocelyn Dupuis, Christine Des Rosiers, Matthieu Ruiz, Julie Hussin

## Abstract

Understanding molecular traits through metabolomics offers an avenue to tailor cardiovascular prevention, diagnosis and treatment strategies more effectively. This study focuses on the application of machine learning (ML) and explainable artificial intelligence (XAI) algorithms to detect discriminant molecular signatures in heart failure (HF). In this study, we aim to uncover metabolites with significant predictive value by analyzing targeted metabolomics data through ML models and XAI methodologies. After robust quality control procedures, we analyzed 55 metabolites from 124 plasma samples, including 53 HF patients and 71 controls, comparing Logistic Regression (Logit) models with Support Vector Machine (SVM) and eXtreme Gradient Boosting (XGB), all achieving high accuracy in predicting group labels: 84.20% (*σ* =5.46), 85.73% (*σ* =6.25), and 84.80% (*σ* =7.84), respectively. Permutation-based variable importance and Local Interpretable Model-agnostic Explanations (LIME) were used for group-level and individual-level explainability, respectively, complemented by H-Friedman statistics for variable interactions, yielding reliable, explainable insights of the ML models. Metabolites well-known for their association with heart failure, such as glucose and cholesterol, but also more recently described association such C18:1 carnitine, were reaffirmed in our analysis. The novel discovery of lignoceric acid (C24:0 fatty acid) as a critical discriminator, was confirmed in a replication cohort, underscoring its potential as a metabolite marker. Furthermore, our study highlights the utility of 2-way variable interaction analysis in unveiling a network of metabolite interactions essential for accurate disease prediction. The results demonstrate our approach’s efficacy in identifying key metabolites and their interactions, illustrating the power of ML and XAI in advancing personalized healthcare solutions.

See *Graphical Abstract*

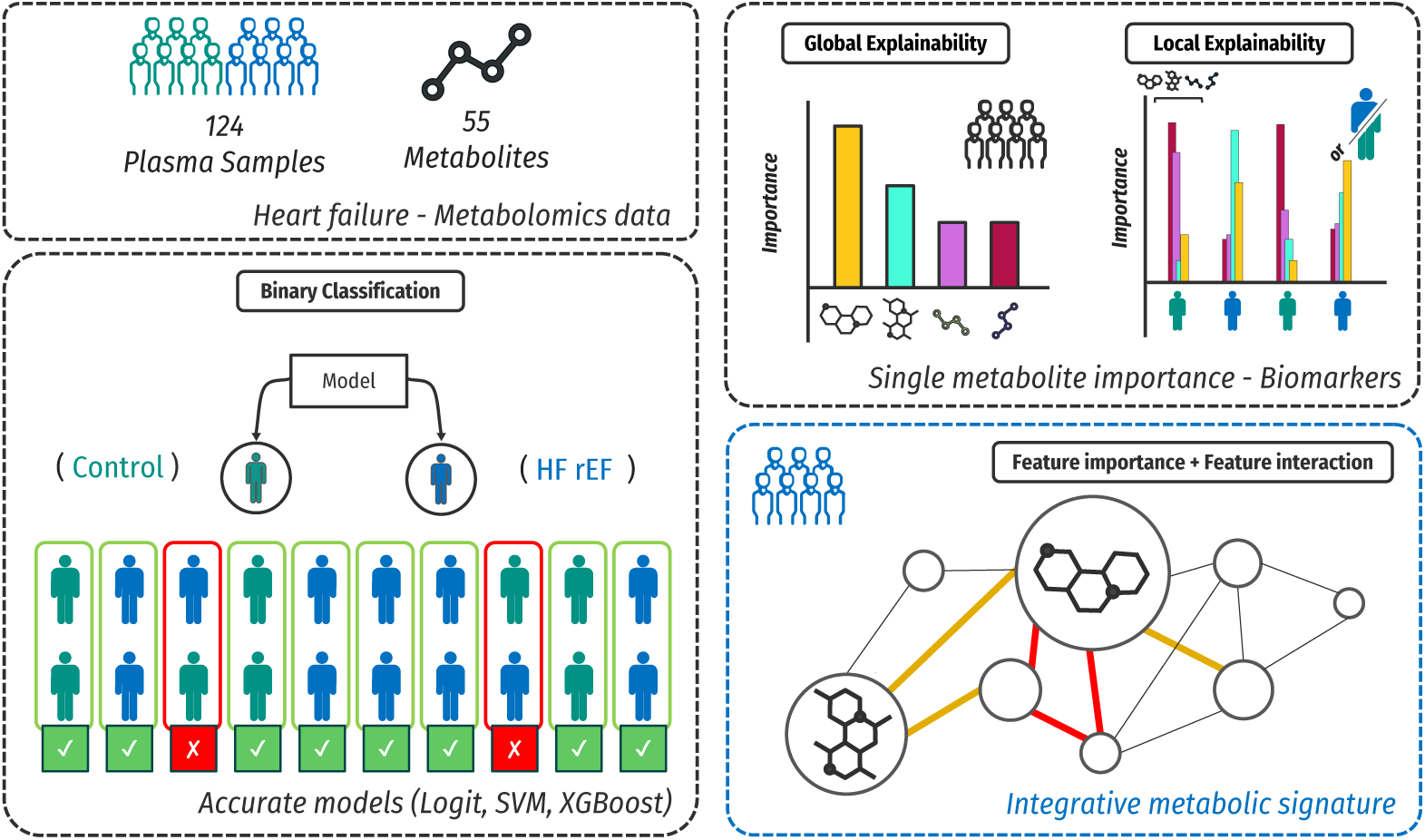

## Introduction

Heart failure (HF) affects 64.3 million individuals worldwide. It is a complex cardiovascular condition characterized by the defective ability of the heart to effectively pump blood throughout the body, unable to meet the body’s metabolic needs [Savarese et al., 2017]. Heart failure can be classified into three subtypes based on the left ventricular ejection fraction (EF) ejection fraction, namely heart failure with preserved ejection fraction (HFpEF *≥* 50%), heart failure with mid-range ejection fraction (HFmrEF, 41-49%), and heart failure with reduced ejection fraction (HFrEF, EF ≤ 40%), each presenting specific associated challenges due to overlapping phenotypes with other pathologies and comorbidities [Ziaeian and Fonarow, 2016]. Independently from the HF types, there is a phenotypic heterogeneity across individuals, due to diverse molecular, and environmental factors they are influenced by. This heterogeneity can be attributed to diverse comorbidities and pathophysiological mechanisms specific to each patient, with HF manifesting as a highly individualized phenotype, thereby accentuating the complexity and challenges in managing the condition effectively. Although therapies for managing heart failure have been developed, and the management of HFrEF has been the most widely characterized, there is still an urgent need to translate it into alternative or complementary therapies. Patients with HFrEF imposes a substantial socio-economic burden on health care systems, and continue to experience high mortality rates. Therefore, it is crucial to better understand disease mechanisms to enable early diagnosis and develop more effective and precise medical strategies [Bloom et al., 2017].

In recent years, omics technologies, including metabolomics protocols, have emerged as a powerful tool to understand the underlying biology of many disease contexts in human, such as cardiovascular diseases [Shah et al., 2010, 2012, Tang et al., 2013], including heart failure [Ahmad et al., 2016, Hunter et al., 2016, Asselin et al., 2014, Ruiz et al., 2017]. Metabolomics aims to comprehensively quantify the small-molecules called metabolites within a biological system, offering insights into metabolic changes associated with the disease states. The identification of new biomarkers specific to the HFrEF patients, can lead to a better understanding of this disease subtype, thus improving precision medicine. These studies typically analyze differential metabolites levels independently, examining them one at a time within linear frameworks, which may not capture the full complexity of the metabolomic-phenotypic relationship as well as the heterogeneity of the disease. The omission of non-linear and interaction effects represents a significant methodological gap for the development of accurate predictive models and the extraction of comprehensive biological signatures.

To take into account the complex nature of metabolic networks underlying heart failure, likely involving non-linear relationships [Mosconi et al., 2008], machine learning (ML) models emerge as a powerful solution to analyze and interpret metabolomics data. Once accurate models are established, explainability algorithms can be leveraged to obtain metabolic profiles associated with heart failure phenotype. Explainability methods determine the importance of each metabolite in the prediction process; it generates an importance score per metabolite, offering insights at both global and local levels. The global explainability scores reflect the overall importance of metabolites across the model for the full set of individuals. Local explainability methods will provide for a specific sample, an importance score of each metabolite, offering potential insights into the underlying disease mechanisms for a specific individual. Local and global importance scores provide valuable insights; they are designed to explain the effects of each variable on the outcome, but variable interactions need to be considered for explaining the remaining effects, here the combinatory effect of several metabolites to improve model classification performance. There are several post-hoc explainability methods, such as partial dependence plot (PDP) [Friedman, 2001], accumulated local effects (ALE)[Apley and Zhu, 2020], permutation variable importance for global explainability[Fisher et al., 2019]. This last method offers several advantages: it is model-agnostic, allowing for comparison across different models, and it relies on a straightforward concept, which is that feature importance is based on the increase in model error when the information from a specific feature is disrupted. Additionnally, local explainability algorithms like individual conditional expectation (ICE) [Goldstein et al., 2015], local interpretable model-agnostic explanations (LIME) [Ribeiro et al., 2016] and Shapley values focus on explaining individual predictions [Lundberg and Lee, 2017]. The LIME algorithm is applicable to any model type, making it a robust and practical choice for our analysis benchmarking multiple classification model types. To detect and quantify interaction effects, interaction scores can be computed per combination of variables, which has the potential to provide a comprehensive understanding of how the predictive model used multiple features in a non-linear way, compared to single-variable importance scores. For instance variable interactions have been measured using various statistical approaches [Loh, 2002, Lou et al., 2013] and decision tree models [Sorokina et al., 2008, Wright et al., 2016, Deng, 2019, Boulesteix et al., 2015]. Recent methods include Friedman’s H-statistic [Friedman and Popescu, 2008], Greenwell’s variance of partial dependency [Greenwell et al., 2020], and Oh’s performance-based measure [Oh, 2019, 2022]. Friedman’s H-statistic is particularly effective in capturing feature interactions and providing scores across various model types, making it a robust option for detailed interaction analysis. To the best of our knowledge, none of the existing explainable artificial intelligence (XAI) methodologies have yet been applied to human metabolomics data within the specific context of heart failure.

In this study, we introduce a comprehensive pipeline that includes robust preprocessing of metabolomics data, exploration of machine learning models for disease classification, application of explainability techniques to ascertain variable importance, and assessment of variable interactions to uncover complex metabolic insights of HFrEF. We applied our methodologies to a dataset from two previously published plasma metabolomics studies [Ruiz et al., 2017, Asselin et al., 2014]. Both studies collectively underscored substantial metabolic dysregulations in heart failure patients. Ruiz et al [Ruiz et al., 2017] demonstrated disruptions in mitochondrial and peroxisomal fatty acid (FA) metabolism through altered acylcarnitine profiles, while Asselin et al [Asselin et al., 2014] identified linoleic acid and HDL-cholesterol as significant determinants of oxidative stress markers. We compared an inherently explainable Logistic Regression model to Support Vector Machine (SVM), a technique chosen for its enhanced classification robustness, and eXtreme Gradient Boosting (XGBoost), chosen for its proficiency in capturing complex non-linear relationships. To understand model decisions, we used permutation-based methods for global explainability and LIME for local insights, which we also compared with Shapley values. We complemented these results by using the H-Friedman statistics to analyze feature interactions. Applying this pipeline to metabolomics data from heart failure studies allowed us to confirm known metabolic markers of heart failure and also to uncover lignoceric acid (C24:0 FA) as a novel discriminant metabolic marker. Our approach demonstrates the strength of integrating machine learning with explainability techniques to analyze the complex metabolomic determinants behind heart failure phenotype.

## Material and Methods

### Discovery dataset preprocessing and quality control for ML input

The discovery dataset in this study was derived from a targeted metabolomics cross-sectional, exploratory analysis within a Montreal Heart Institute (MHI) cohort focused on investigating blood circulating metabolites in HFrEF patients. The specific inclusion and exclusion criteria and methodological details for assessing plasma metabolites have been described previously [Asselin et al., 2014, Ruiz et al., 2017]. Our dataset comprises 132 plasma samples, categorized into controls (n = 72) and HFrEF patients (n = 60) for which 71 metabolites were measured. When applicable, metabolite names will specify carbon atoms and number of unsaturation, separated by a colon (”:”)(see [Ruiz et al., 2017]). Samples with more than 25% missing values were excluded (n=8), and, for the remaining samples, metabolites with more than 10% missing values were excluded (n=13) (Supplementary Table S1). The following data preparation steps were done using the online R version of MetaboAnalyst 5.0 (www.metaboanalyst.ca, last accessed on 01-24-2023) [Xia et al., 2009]. Remaining missing values were imputed using the K-Nearest Neighbors algorithm. Near-constant metabolites with low interquartile range (IQR), likely to be non-informative variables, were removed (n=3). Our final dataset contains 55 metabolites measured in 124 samples. To reduce biases due to highly variable metabolites’ range, raw concentrations underwent auto-scaling and log transformation. Additionally, to control for confounding variables such as age and sex, we corrected these standardized concentrations by employing the residuals from a linear model that fitted each metabolite level against these covariates. We conducted Principal Component Analysis (PCA) on this pre-processed dataset to identify any remaining quality issue such as outliers or batch effects, that could potentially affect the reliability of subsequent machine learning experiments.

### Replication cohort

To validate our marker discovery, we utilized a replication cohort derived from a comprehensive metabolomics dataset. The cohort was part of a larger clinical study aimed at investigating metabolic profiles and their association with left heart disease (LHD) with or without pulmonary hypertension (PH). The OILY cohort, for Evaluation of the Metabolic Pr<u>o</u>file as an Indicator for Group II Pulmonary Hypertension, consists of individuals selected based on predefined inclusion and exclusion criteria to ensure the robustness and reproducibility of the results. Briefly, the inclusion criteria for subjects were: males and females aged 18 years or older, referred for a medically required left and right heart catheterization for their cardiac condition, with LHD of all etiologies at risk of PH according to the World PH Symposium, must be stable without hospitalization in the last 3 months, females must be menopausal or surgically sterilized; otherwise, they must have a negative pregnancy test at the time of heart catheterization. Exclusion criteria were: subjects with known PH of other groups or at risk of PH not due to LHD, and pregnant women. For comparison, age- and sex-matched healthy subjects from the MHI biobank in Quebec were used. The study was conducted following the Declaration of Helsinki and approved by the Montreal Heart Institute Ethics Committee (ICM 2019-2538). Informed consent was obtained from all participants. Data privacy and confidentiality were maintained according to institutional and regulatory guidelines. An arterial blood sample were collected from participants via right radial artery cannulation in a subset of patients. For plasma preparation, blood were collected in EDTA tubes (BD Vacutainer) and immediately placed on ice for 30 min before centrifugation at 2,000 g for 15 min. The resulting plasma were then aliquoted into Sarstedt microtubes, frozen on dry ice before storage at -80°C until analysis. Plasma samples were processed for quantitative profiling of fatty acids using gas chromatography-mass spectrometry (GC-MS) following a previously described method [Pinçon et al., 2019]. Briefly, plasma samples (100 *μL*) were incubated overnight at 4 °C in a chloroform/methanol solution (2:1) containing 0.004% butylated hydroxytoluene (BHT). The mixture was filtered through gauze, dried under nitrogen gas, and re-suspended in hexane/chloroform/methanol (95:3:2). After adding internal standards, all samples were dried under nitrogen gas and re-suspended in hexane/methanol (1:4) containing 0.004% BHT. C24:0 fatty acids were analyzed as their methyl esters following direct trans-esterification with acetyl chloride/methanol. This was performed on a 7890B gas chromatograph coupled to a 5977A Mass Selective Detector (Agilent Technologies, Santa Clara, CA, USA) equipped with a capillary column (J&W Select FAME CP7420; 100 m × 250 μm inner diameter; Agilent Technologies, Santa Clara, CA, USA). The system operated in positive chemical ionization mode using ammonia as the reagent gas. Samples were analyzed under the following conditions: injection at 270 °C in split mode using high-purity helium as the carrier gas, with a temperature gradient of 190 °C for 25 minutes, increased by 1.5 °C/min up to 236 °C. C24:0 FA were detected as their [M+NH3]+ ions by selective ion monitoring, and their concentrations (in *μM*) were calculated using standard curves and the labeled D47-lignoceric acid standard. To ensure the reproducibility of our metabolite marker discovery, we utilized a subset derived from the comprehensive OILY metabolomics dataset. This subset of the replication cohort included 71 controls, 31 heart failure samples, and specifically 14 HFrEF samples (LVEF ≤ 40%) for which C24:0 FA levels were measured (Supplementary Table S2).

### Classification algorithms

We used supervised machine learning algorithms to classify individuals into two groups: those diagnosed with HFrEF and controls, using three distinct algorithms : Logistic Regression (Logit), Support Vector Machines (SVM), and eXtreme Gradient Boosting (XGBoost). To ensure the robustness of our models with small sample size, we evaluated them through a 5-fold cross-validation procedure using “caret” package in R (version 6.0-93, [Kuhn, 2022]). This involved generating training sets through stratified random splits of the data, and performing a grid search for hyperparameters tuning. All models were utilized with default hyperparameters. If any parameters were tuned, the details can be found in Supplementary Table S5 and Supplementary Table S6. Logit models will serve as a baseline to better understand the complexity of our task. Prior to fitting the Logit model, our baseline model, variable selection was performed to minimize the number of variables, thereby preventing multicollinearity and overfitting. We first removed variables with Spearman correlation coefficient above 0.8 and then employed the F-score metric from the “FSinR” package in R (version 1.0.8, [Aragón-Royón et al., 2020]), ultimately retaining 10 metabolites for the Logit model. For the SVM classifier, we opted for the radial basis function kernel, due to its superior accuracy and its capability to effectively model non-linear relationships among metabolite variables, using “e1071” package in R (version 1.7-13, [Meyer et al., 2023]). We fitted the XGBoost (Extreme Gradient Boosting) models using the “xgboost” package in R (version 1.7.3.1, [Chen et al., 2023]). The models’ performances were evaluated using classical metrics such as accuracy, specificity and sensitivity, as well as the balanced accuracy metric, providing a comprehensive assessment of a model’s ability to correctly classify samples while addressing class imbalance.

### Explainability and variable interactions

To assess the global explainability of our models, we used the “DALEX” package in R (version 2.4.3, Biecek [2018]), implementing a permutation-based method. This method involves systematically omitting one or more variables from the data input, then re-evaluating the model’s performance to see how it changes without those variables. The type of importance was specified as the “ratio” method, to focus on the proportional change in the model’s prediction error. For quantifying the prediction error, we chose the cross-entropy (or “log loss”) loss function, which provides a measure of the mean of magnitude of the error, over 500 iterations. For each sample, we assessed the local explainability by using the LIME algorithm using the “lime” package in R (version 0.5.3, [Hvitfeldt et al., 2022]) with default parameters. A variable weight is computed for each metabolite and each sample, with values that can be either negative or positive (from -1 to 1), either contradicting the probability or supporting the probability to be classified as heart failure, respectively. We compared the obtained LIME weight with SHAP values using the “SHAPforxgboost” package in R (version 0.1.3,[Liu and Just, 2023]). Variable interactions were analyzed using the H-Friedman statistics from the “iml” package in R (version 0.11.1, [Molnar et al., 2018]) with default parameters, to quantify how combinations of variables non-additively improve prediction performance in our heart failure classification models. All methods were utilized with default hyperparameters, except where noted in the previous paragraph.

### Statistical analysis

All statistical analyses were conducted using R (version 4.1.2). The normality of the data was assessed using the Shapiro-Wilk test. For the comparison of two independent groups, we employed the Wilcoxon rank-sum test, also known as the Mann-Whitney U test as non normality was assessed by Shapiro test. All statistical tests were two-tailed, and the significance level was set at 0.05. Results were presented with corresponding raw p-values.

### Data and code availability

The discovery cohort data underlying this article are available in Mendeley Data at https://data.mendeley.com/, and can be accessed with DOI: 10.17632/57fsdkv6gh.1. The code developed in this study have been made available to ensure transparency and reproducibility of our findings and can be accessed at the following public repository: https://github.com/HussinLab/ML-XAI-HF. All analyses were conducted using R version 4.1.2, and the codebase includes scripts for machine learning model implementation and explainability analysis.

## Results

To explore the potential of analysing metabolomics data with machine learning models plus post-hoc explainability methods, we developed a pipeline, presented in Figure 1. Our curated dataset contains 124 plasma samples (53 patients and 71 control samples), with measurements of 55 metabolites. These include common metabolic markers such as cholesterol, triglycerides, glucose, insulin and glycerol, as well as fatty acids (total fatty acids free and bound to phospholipids and triglycerides), organic acids (ketone bodies, lactate, pyruvate and Krebs cycle intermediates), and acylcarnitines, which serve as indicators of altered fatty acid utilization and oxidation. Our analyses focused on four main aspects : binary classification of control and HFrEF samples using Logit, SVM, XGBoost models, then global explainability using a permutation-based method, and local explainability via LIME, and finally variable interactions through the H-Friedman statistic.

**Fig. 1.**
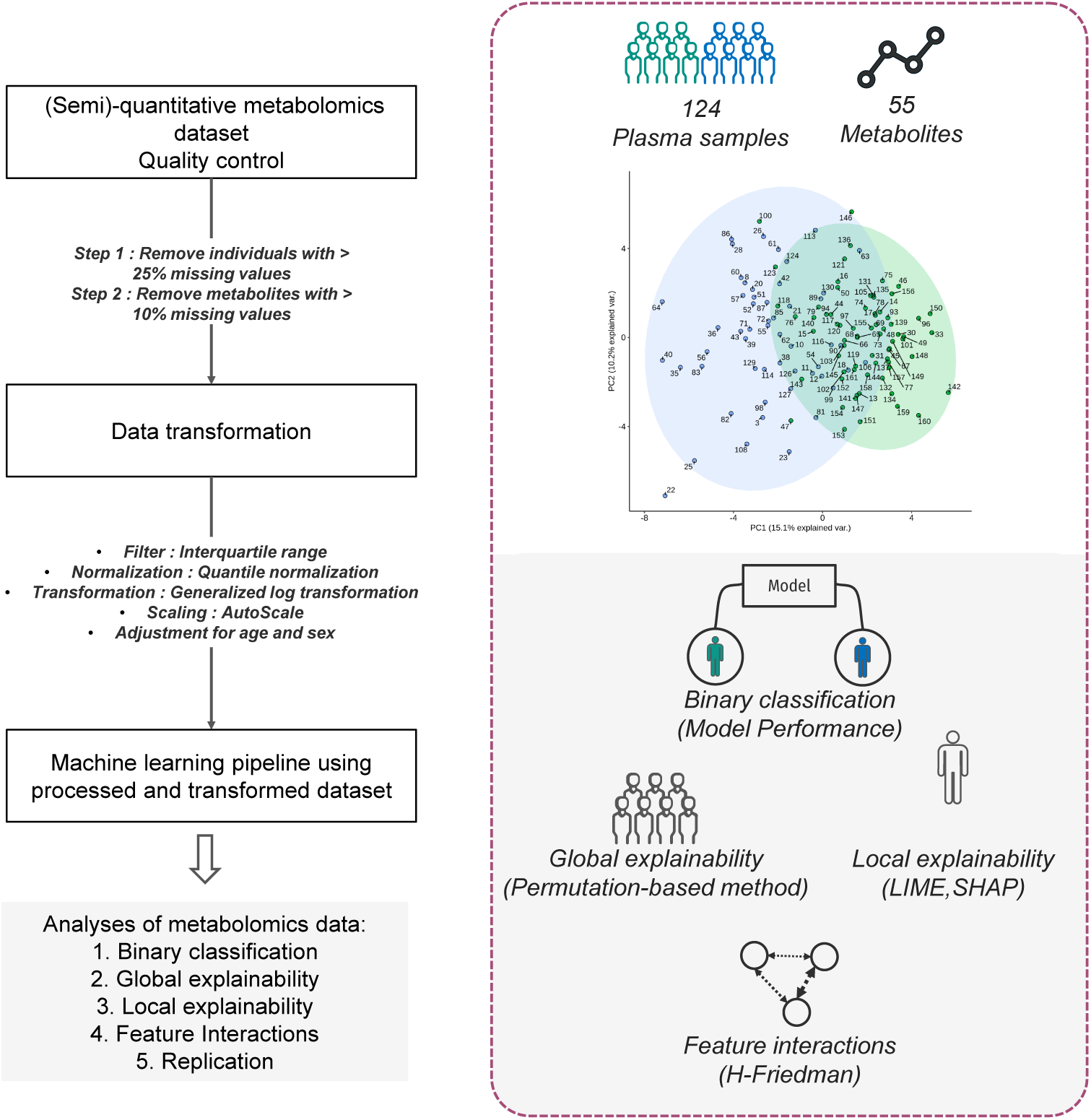
Overview of the study. On the left panel, we show the main steps of the pipeline used to obtain different classification models discriminating between control and samples with heart failure. Boxes indicate input and outputs of processing steps described on arrows. The right panel shows the details of the input data; 124 plasma samples for 55 metabolites and the four main assessments made from the input data

### Distinguishing heart failure patients from controls

To perform quality control for the machine learning input matrix, we conducted a PCA analysis, which reveals a clear distinction between the control and HFrEF classes, with a gradual separation along the PC1 axis (Supplementary Figure S1) without any anomalous clusters, suggesting the dataset’s suitability for fitting our classification models to discriminate controls and heart failure samples.

We next trained different models and assessed their performance. The machine learning performance results for classifying patients and controls based on metabolite measurements for three algorithms, evaluated using cross-validation (see *Material and Methods*) are summarized in Table 1. Our analysis revealed distinct patterns in the performance of the machine learning algorithms by using a 5-cross validation scheme. The logistic regression trained on 10 variables in most folds (see Supplementary Table S4), which served as baseline, captured linear relationships exclusively, demonstrated satisfactory performance, with an mean accuracy of 84.20% (*σ* =5.46). However, the SVM and XGBoost models consistently outperformed it, showcasing mean accuracies of 85.73% (*σ* =6.25) and 84.8% (*σ* =7.84), respectively. Importantly, while comparing SVM and XGBoost models, we observed greater stability in the performance metrics of XGBoost, with four metrics scoring above 80% (Table 1). When comparing different seeds, one (seed 124) demonstrated higher stability across the five folds, leading to its arbitrary selection for subsequent analysis (Supplementary Table S3). The list of wrongly classified samples can be found in Supplementary Figure S4 and also highlighted in the PCA Supplementary Figure S1, with several samples that were consistently misclassified in the XGBoost models (IDs: 154, 143, 116, 113, 106, 99, 97, 76, 63, 47). To test the utility of obtaining explanations from these models in understanding the phenotype within the context of metabolomics, we opted for subsequent explainability analysis of the SVM and XGBoost models trained.

**Table 1.** Overview of model performances, for the 5 cross-validation schemes, across 3 different seeds.

| Performance Metric Name | Logit | SVM | XGBoost |
| --- | --- | --- | --- |
| Specificity | 86.47 - 9.61 | 91.20 - 5.58 | 82.00 - 6.98 |
| Sensitivity | 81.33 - 10.70 | 78.47 - 9.87 | 86.87 - 9.80 |
| Balanced Accuracy | 83.80 - 5.42 | 84.67 - 6.34 | 84.33 - 7.49 |
| Accuracy | 84.20 - 5.46 | 85.73 - 6.25 | 84.80 - 7.84 |

### Explaining reliable models identify well-known and novel metabolite markers

The evaluation of the SVM and XGBoost models using a 5-fold cross-validation scheme demonstrated high performance across all folds illustrating the models’ reliability, regardless of the specific samples included in each fold (see *Distinguishing heart failure patients from controls*). Since each fold may produce fold-specific explainability profiles, our objective was to obtain a single model explainability to avoid potential batch effects arising from fold sample composition. To achieve a consistent global and local explainability, we retrained a single model for both SVM and XGBoost algorithms using the entire dataset of 124 samples and 55 metabolites (*”full model”*) and validated that the results align with the fold-specific models (see Supplementary content, Supplementary Figure S2).

We next computed dropout loss to gauge the impact of dropping out metabolites from our models, and understand their global importance and influence on the model’s predictions. The change in model performance, measured as an increased error rate, indicates the importance of the dropped variables in making predictions, allowing us to identify the top 10 global importance scores for SVM and XGBoost models (Figure 2). We observe distinct behaviors between the SVM and XGBoost models regarding global importance scores. The SVM exhibits relatively lower variation when applied to the permuted matrix, as illustrated by the arithmetic mean of ratio values slightly *>* 1. In contrast, the XGBoost model demonstrates greater volatility, as indicated by the taller bars and the mean ratio values *>* 1, with a maximum mean ratio value of 2.80. This illustrates the XGBoost capacity to prioritize a few metabolites for making predictions. Both models identified a common set of critical metabolites, including glucose, various carnitines, long-chain fatty acids, triglycerides, and cholesterol, indicating robust global explainability. This overlap in variable importance between theses two algorithms highlights the robustness of global explainability. Specifically, glucose was the most discriminant metabolite for the SVM, ranking third in the XGBoost model, while C24:0 FA was the most discriminant for the XGBoost model, ranking third in the SVM model, underscoring their potential roles in predicting heart failure with reduced ejection fraction.

**Fig. 2.**
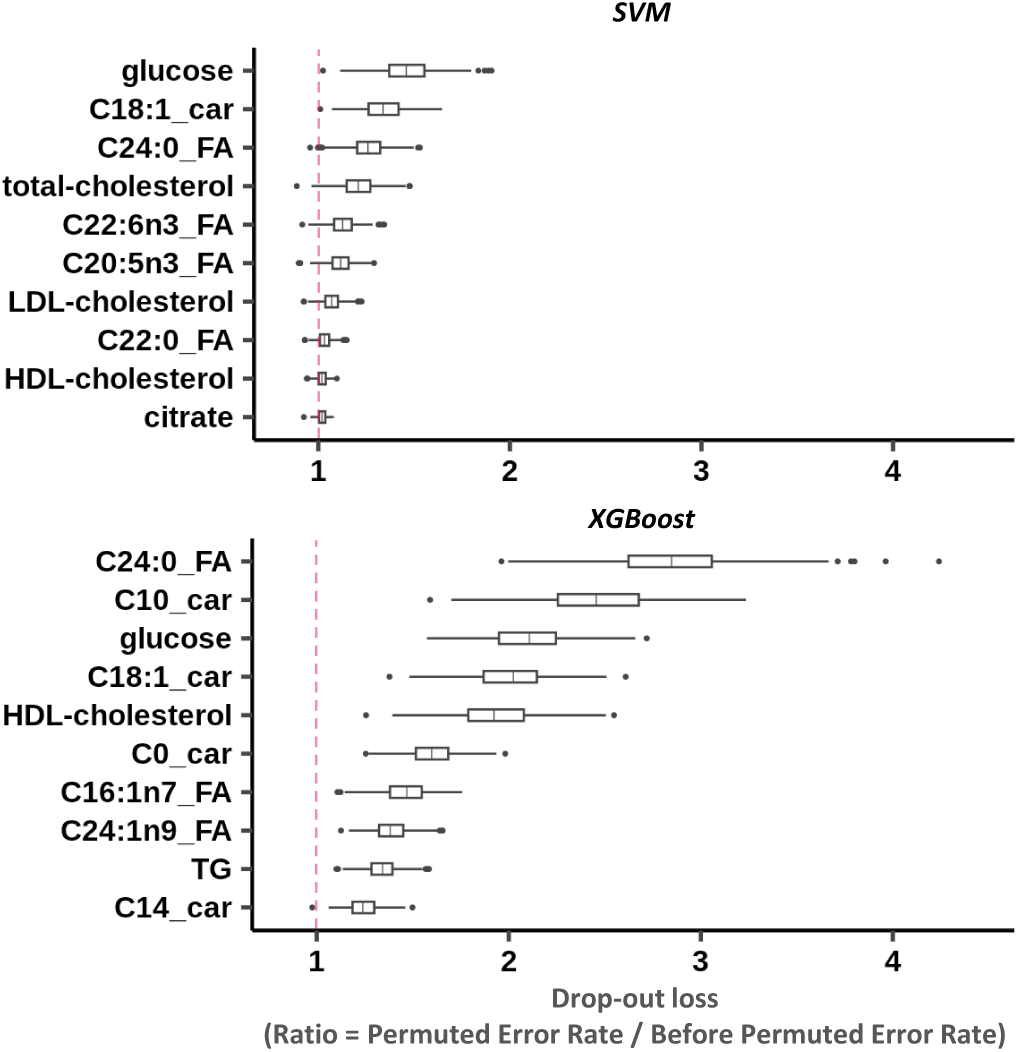
Global explainability by using a permutation-based methodology. Boxplots of the top 10 most important variables at the global scale for the SVM and XGBoost models. Variable importance is determined by calculating the increase in the model’s prediction error after permuting the variable. Here, we permuted each variable 500 times. The error metric used is the cross-entropy loss function. The x-axis represents the ratio of the prediction error of the model with the permuted variable matrix to the prediction error of the baseline model. The y-axis shows the top 10 metabolites. Boxplots display the ratio values for all different permutations. Boxplot components: Central line indicates the median, the box captures the upper and lower quartiles, bars extend to 1.5 times the interquartile range, points outside these bars represent outliers. Vertical dotted red lines indicate the value at which no difference is observed after permuting the variable. Abbreviations - car = Carnitine, FA = Fatty Acid, HDL = High-Density Lipoprotein, LDL = Low-Density Lipoprotein, SVM = Support-Vector Machine, TG = Triglycerides, XGBoost = eXtreme Gradient Boosting.

To address the inherent heterogeneity of the HFrEF phenotype, we use local explainability approaches to enhance our understanding of how individual metabolites uniquely influence model predictions across different samples (Figure 3). Using the LIME methodology (see *Material and Methods*) for each model and each metabolite, we obtain a weight value that represents an importance score indicating whether the sample’s metabolite level supports the probability of being classified in the heart failure class (indicated by positive values) or contradicts it (indicated by negative values). These importance scores can be presented in a heatmap of the local explainability for each of the 124 samples (Figure 3). The SVM model (Figure 3-A) shows a greater level of variability in local importance scores across the different metabolites and samples compared to the XGBoost model (Figure 3-B, Supplementary Figure S3). While the results allow for the assessment of individual metabolites’ contributions to the prediction process, this variability indicates a broader distribution of importance scores among metabolites, complicating the identification of specific important biomarkers. In contrast, the XGBoost model’s heatmap displays only a restricted number of densely colored columns, indicating its ability to prioritize a specific subset of metabolites for the prediction task. Here, by using heatmap representations, we observed that XGBoost LIME profiles are more distinct in comparison to those of SVM, as illustrated by the use of a restrained subset of metabolites. It is difficult to pinpoint a subset of metabolite for the SVM model, but we can cleary see that C24:0 FA is the most discriminant one for the XGBoost model. We compared LIME and SHAP values and determined that both methods are consistent in their observations for all values, but also specifically for the C24:0 FA (see Explainability and variable interactions, Supplementary Figure S6).

**Fig. 3.**
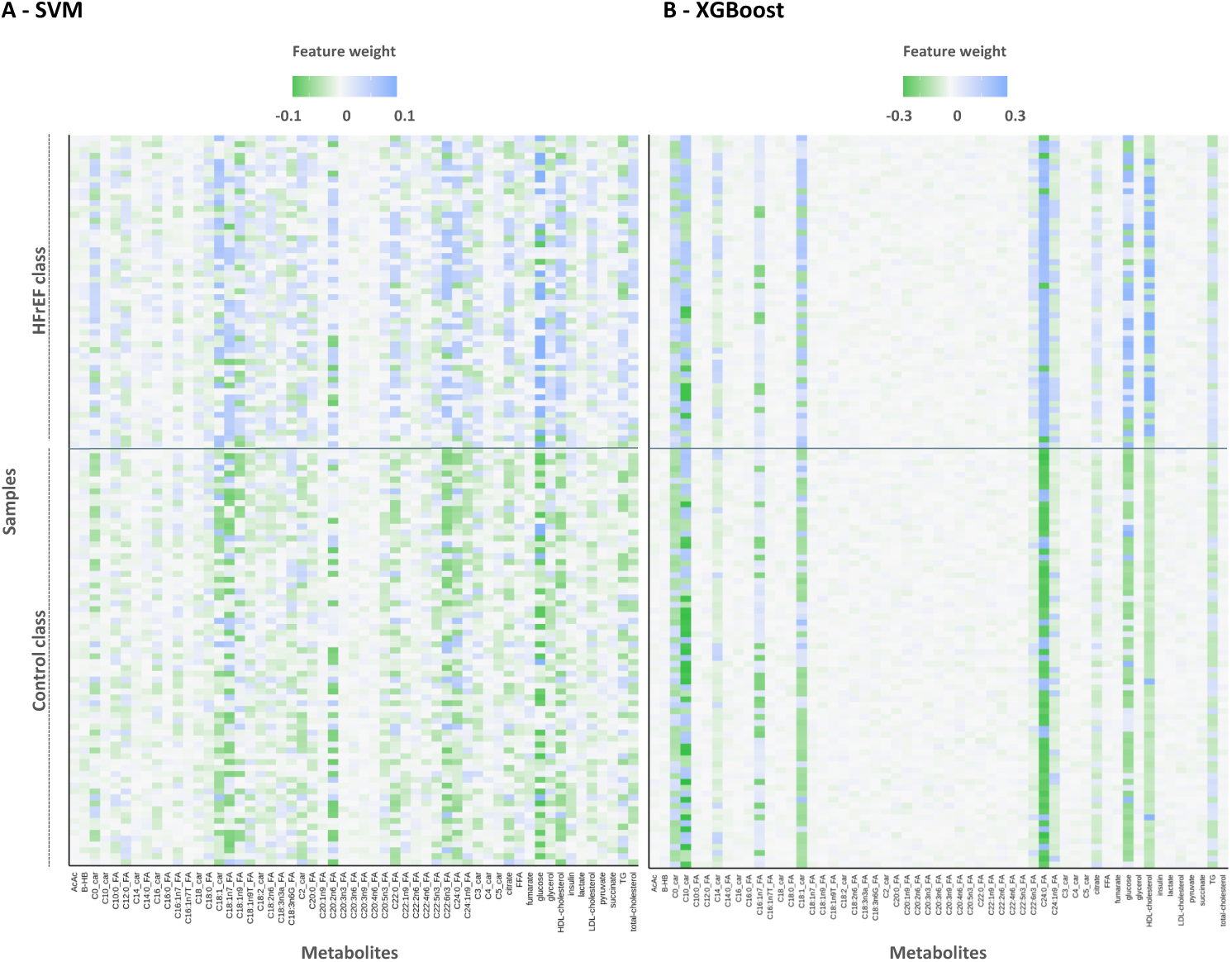
Local explainability using LIME for the SVM and XGBoost model. Heatmaps of the variable weights for 55 metabolites, for the 124 samples. Metabolites are on the x-axis, and the y-axis denotes individual cases. Variable weights signify the contribution of each variable to support the probability of heart failure, with positive values depicted in blue. Variable weights contradicting the probability of heart failure, with negative values, are represented in green. A densely colored square in the heatmap signifies the importance of a particular metabolite for a specific sample. Abbreviations - AcAc = Acetoacetate, BHB = Beta-hydroxybutyrate, car = carnitine, FA = Fatty Acid, FFA = Free Fatty Acids, HFrEF = Heart Failure with Reduced Ejection Fraction, HDL = High-Density Lipoprotein, LDL = Low-Density Lipoprotein, TG = Triglycerides.

### Identification of interacting metabolites

The heatmap representation (Figure 3) also suggests that the predicted phenotype may involve interactions, as individual LIME profiles do not rely on a single metabolite but rather require information from multiple metabolites to accurately predict the probability of heart failure, as observed across various samples. We aim to reveal potential interaction effects by exploring 2-way interactions between features used within the different machine learning models we trained. The H-Friedman statistic offers a statistical score of interaction (see *Material and Methods*). According to this metric, our results indicate that most metabolites are not involved in any interactions, with 43 metabolites out of 55 with an interaction score of zero (78.18%) (Figure 4-A). For a subset of 12 metabolites, however, their H-Friedman scores have non-zero values, confirming that there is a small proportion of metabolites (21.82%) involved in interactions and that together they are improving the model’s predictions (Figure 4-A). Notably, the metabolites involving interactions are the top 10 most important metabolites derived from our global and local explainability analyses (Figure 2, 3), in addition to two that were not defined as main contributors to accurate predictions, namely citrate and C22:6n3 fatty acid. Among the pairs of interacting metabolites, several have interaction scores particularly larger than others, due to their high influence on the binary classification model’s performance, as supported by the H-Friedman statistic (Figure 4-B). Notably, C24:0 FA, C18:1 carnitine, glucose, and HDL-cholesterol stood out as both individually important and central to various interactions.

**Fig. 4.**
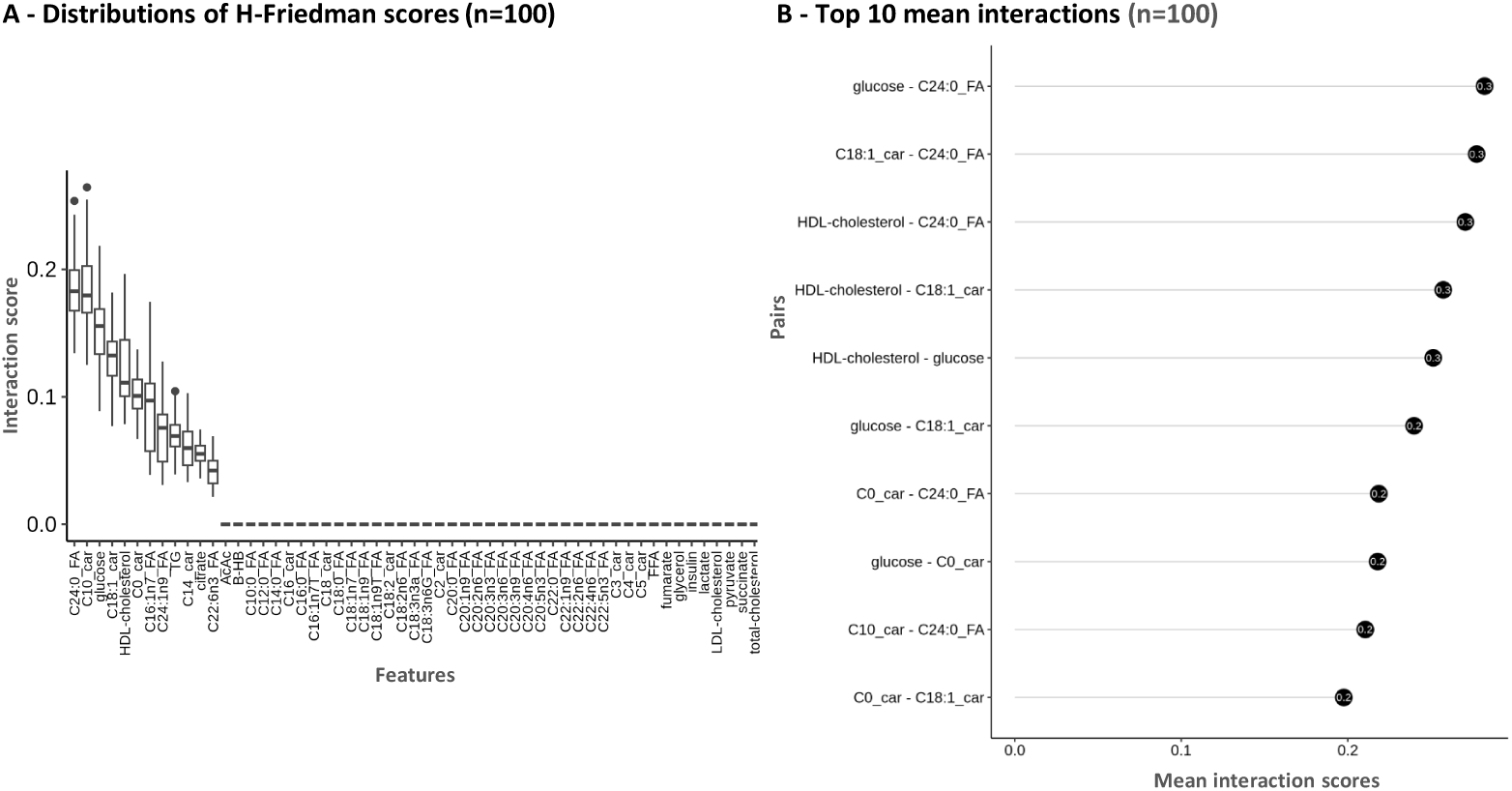
Variable interactions of the XGBoost model. A - Distribution of all 2-ways interaction scores for the 55 variables (n = 2970) of the 124 samples for 100 permutations. B - Top 10 interaction strengths based on mean of interaction scores for each pair. Abbreviations - AcAc = Acetoacetate, BHB = Beta-hydroxybutyrate, car = carnitine, FA = Fatty Acid, FFA = Free Fatty Acids, HFrEF = Heart Failure with Reduced Ejection Fraction, HDL = High-Density Lipoprotein, LDL = Low-Density Lipoprotein, TG = Triglycerides.

### Most important metabolites are also used for misclassified samples

The classification task involving three different seeds and algorithms resulted in some misclassified samples (Supplementary Figure S4-A/B). LIME explanations are valuable for performing error analysis on individual predictions. Understanding the reasons behind these misclassifications is important for addressing dataset limitations and task complexity. This analysis aims to identify variables contributing to misclassifications and verify potential model biases towards the use of non-informative variables. Additionally, we can verify if these samples have overall metabolic profiles that diverge greatly from those expected for their true labels, such as a sample with a heart failure label exhibiting control-like values. Focusing on the XGBoost models, ten samples were consistently misclassified (IDs: 154, 143, 116, 113, 106, 99, 97, 76, 63, 47), highlighted in the PCA analysis shown in Supplementary Figure S1. Overall, the most important metabolites for misclassifying these samples are consistent with those important for the correctly classified samples, namely C24:0 FA, C18:1 carnitine, and glucose (Supplementary Figure S5). We analyzed the worst predictions, those with the largest prediction errors for control and HFrEF cases, referred to as false positive and false negative examples, with probability outputs of 1.00 and 0.03, respectively (Supplementary Figure S4-C/D), showing the model’s confidence in these predictions given as shown by these extreme probabilities. In these cases, we can observe that we have a subset of the most important metabolites in the top variables which remains consistent with the global explanation, but their weights are reversed due to the metabolite values. Overall, this is illustrating the difficulty of the classification task for these samples and most likely a dataset limitation.

### Combining explainability and variables interactions

Given the interactions uncovered between important features from the XGboost model, we visualized the metabolic signature of the disease phenotype using network graph visualization, which combines feature importance from the global explanations (Figure 2) with interaction scores (Figure 4). The interactions among all variables in the XGBoost model form a subnetwork within the broader metabolite landscape, indicating that the model captures numerous relationships within the data to make accurate predictions (Figure 5-A). We prioritized a subnetwork by focusing on the top 12 variables with the highest interaction scores to understand their implications for the binary classification task (Figure 5-B). From this, we observe that the most central metabolites are C24:0 FA, C18:1 carnitine, glucose, and HDL-cholesterol (Figure 5-B). Additionally, some key interactions involve minor variables such as citrate and free carnitine (C0). These findings offer a detailed view of the interaction landscape within the model’s performance, enhancing our understanding of the classification task, thus the disease phenotype.

**Fig. 5.**
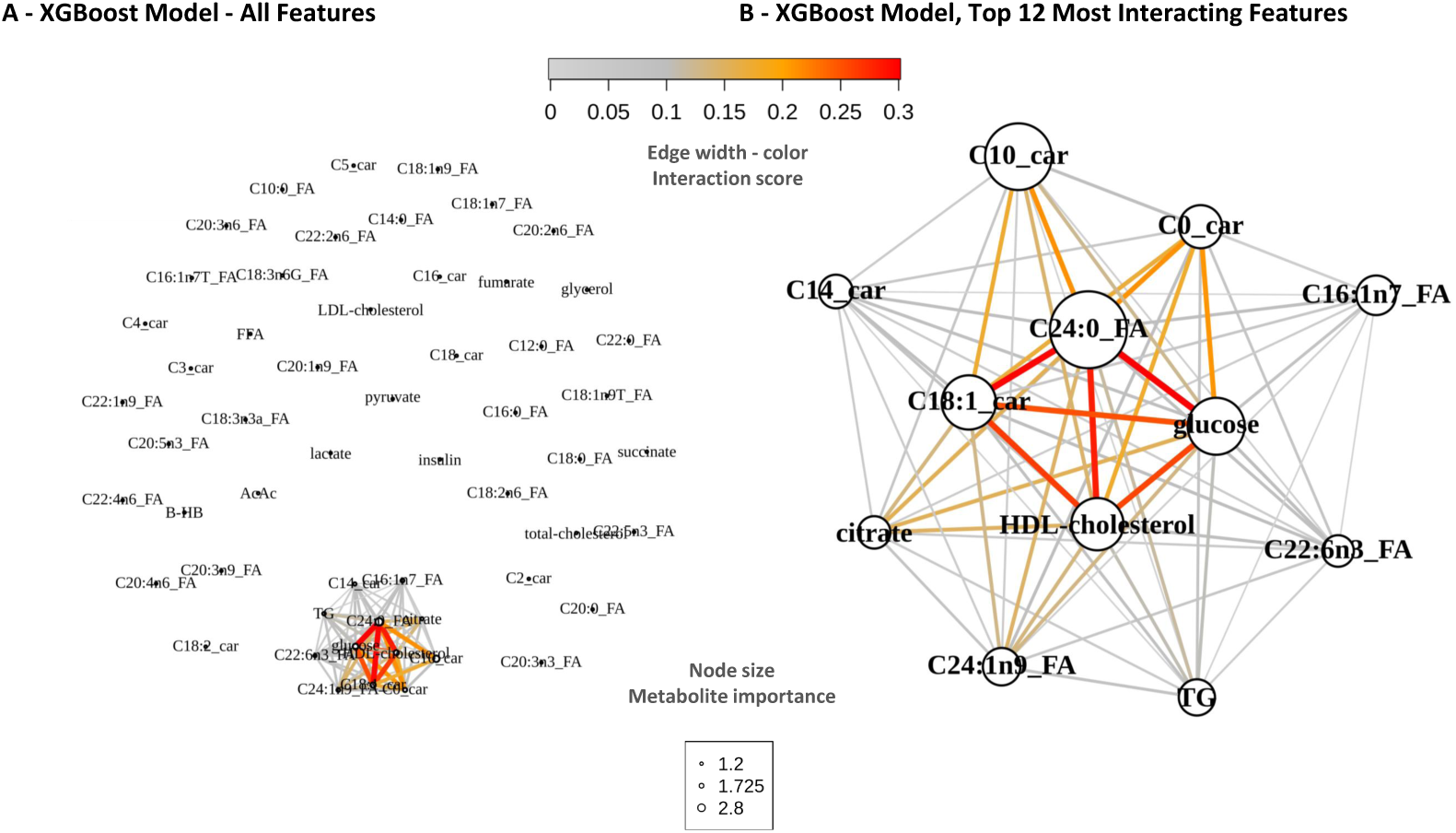
Explainability network of the XGBoost model, combined with interaction scores. Network representing variable as nodes from explainability and interactions score as edge from the H-Friedman statistics. A - All variables used in the XGBoost model are displayed, edges are colored for positive interaction scores. B - A filtered network highlighting metabolites with overall interaction strengths above 0, edges showing positive interaction scores, nodes representing importance of each metabolite based on the global explainability drop-out loss value from Figure 2. Abbreviations - AcAc = Acetoacetate, BHB = Beta-hydroxybutyrate, car = carnitine, FA = Fatty Acid, FFA = Free Fatty Acids, HFrEF = Heart Failure with Reduced Ejection Fraction, HDL = High-Density Lipoprotein, LDL = Low-Density Lipoprotein, TG = Triglycerides,XGBoost = eXtreme Gradient Boosting.

### Replication of the lignoceric acid metabolite marker (C24:0 FA) association with heart failure with reduced ejection fraction

Our workflow allowed us to identify a novel metabolite, the lignoceric acid metabolite marker (C24:0 FA), of potential importance to discriminate between HFrEF cases and controls. We aimed to replicate this result in independent heart failure samples from a replication cohort and using a more robust, quantitative GC-MS method for profiling of plasma total fatty acids free and bound to phospholipids and triglycerides. We utilized a replication cohort (see Discovery dataset preprocessing and quality control for ML input) containing 71 control samples, 31 heart failure patients samples, which includes a subset of 14 HFrEF samples. To test whether the C24:0 FA was discriminatory in this replication cohort, we performed a Wilcoxon test comparing metabolite levels between the groups (Figure 6). The C24:0 FA exhibits lower concentration values in the HF group compared to control (Wilcoxon test, p-value = 1.0 *×* 10*^−^*^13^, Figure 6-A), which is also the case when only HFrEF patients are considered (Wilcoxon test, p-value = 4.9*×*10*^−^*^8^, Figure 6-B), thereby confirming the strong association between C24:0 FA and heart failure uncovered through global and local explainability analyses of our predictive models in the discovery cohort. To evaluate the clinical relevance of the metabolite marker C24:0 FA, we analyzed its correlation with two different disease markers within the HF and HFrEF groups. A significant positive correlation is observed between C24:0 FA concentrations and estimated glomerular filtration rate (eGFR) values in HF samples overall (Pearson R = 0.41, p-value = 0.02) and specifically in the HFrEF subgroup (R = 0.63, p-value = 0.01) (Figure 6-C/D). Additionally, there is a negative correlation between C24:0 FA concentrations and NT-proBNP levels for both HF and HFrEF samples (Pearson R = -0.19, p-value = 0.31, R = -0.49, p-value = 0.07, respectively) (Figure 6-C/D). These findings suggest that C24:0 FA concentration may also serve as a marker of disease severity.

**Fig. 6.**
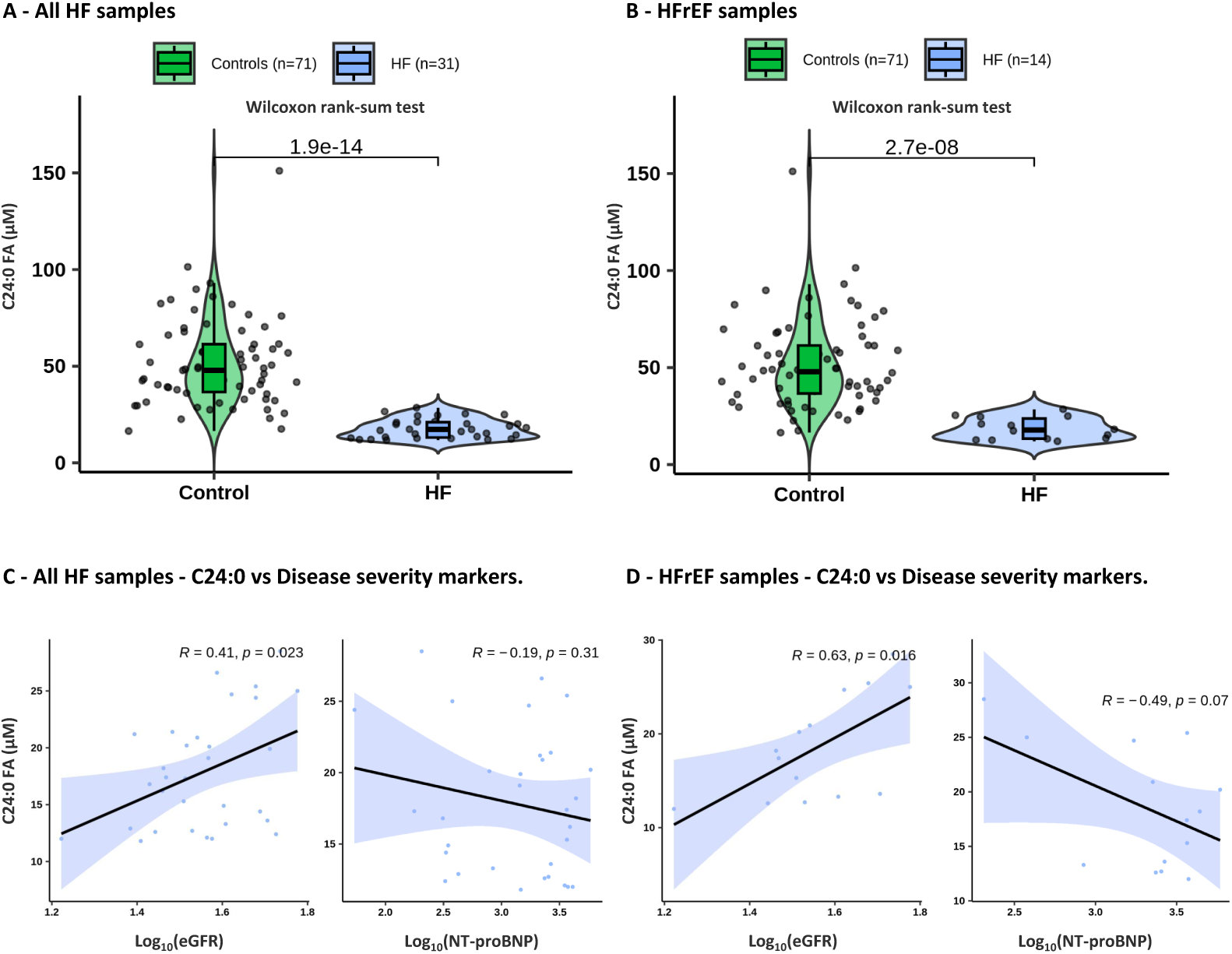
Replication of C24:0 FA using replication cohort. Violin plots of the C24:0 FA concentration (in *μM*), distribution test using Wilcox.test comparing controls and HF samples. A - Control samples (n=71) vs HF samples (n=31) without any filter related the left ventricular ejection fraction B - Control samples (n=71) vs HFrEF samples only (n=14) by using left ventricular ejection fraction (LVEF) value as filter (LVEF ≤ 40%). Correlation plots between C24:0 FA concentrations and log10 heart failure disease markers values. Left : eGFR values, Right : NT-proBNP. C - Correlation using the HF samples (n=31). D - Correlation using the HFrEF samples only (n=14). Abbreviations - eGFR : estimated Glomerular Filtration Rate, FA : Fatty Acid, NT-proBNP : N-terminal pro B-type Natriuretic Peptide.

## Discussion

Physiopathological phenotypes such as HF result in significant costs in terms of human lives and financial resources. To address these challenges, a deeper understanding of the underlying molecular traits is essential. Metabolomics offers an avenue to tailor prevention, diagnosis, and treatment strategies more effectively for various such phenotypes. In this study, we developed a computational pipeline that uses explainable machine learning for the effective analysis of metabolomics data by considering non-linear relationships to capture the heterogeneous complexity of such phenotypes. We used two distinct machine learning algorithms, SVM and XGBoost, compared to baseline logistic regression, to classify control and HFrEF samples based on 55 metabolites’ concentration. Using a dataset of 124 plasma samples, we evaluated model performance, explainability, and identified important metabolites associated with HFrEF. Our analysis reaffirmed known HF-associated metabolites, such as glucose, cholesterol, and C18:1 carnitine. Indeed, our findings align with previous research where notable alterations have been found in glucose [Tran and Wang, 2019, Nielson and Lange, 2005] and cholesterol levels [Rauchhaus et al., 2003, Freitas et al., 2009] in HF patients. Long-chain acylcarnitine have been described to be significantly associated with HF, and they can distinguish between HFrEF and HFpEF patients [Hunter et al., 2016]. Moreover, elevated C16 and C18:1 acylcarnitines in end-stage heart failure patients are linked to higher readmission and mortality risks [Ahmad et al., 2016], but also to severity marker of disease (NT-proBNP) Ruiz et al. [2017], Tremblay-Gravel et al. [2021]. Notably, we identified lignoceric acid (C24:0 FA) as a novel important discriminator, a result that we replicated in an independent cohort with metabolites measured with a different metabolomics protocol. Additionally, two-way variable interaction analysis revealed a network of interacting metabolites, leading to the identification of a more complete/accurate/biology-like integrative heart failure signature encompassing key metabolites and their interactions. These results demonstrate the efficacy of our approach and the power of machine learning and XAI in advancing personalized healthcare solutions.

The potential of machine learning models for classification is significant in enhancing diagnostic accuracy across various medical conditions, including in heart disease [Ibrahim and Abdulazeez, 2021, Safdar et al., 2018]. These algorithms can identify complex patterns and relationships in data that traditional methods might miss. When combined with metabolomics, which provides a comprehensive snapshot of metabolic changes associated with diseases, these models can be particularly powerful. Metabolomics has already shown utility in diagnosing heart failure, providing a metabolic signature that can aid in early and accurate detection [Cheng et al., 2015]. In our work, we demonstrated the utility of combining metabolomics and machine learning to obtain accurate classification models distinguishing HFrEF samples from control, achieving prediction accuracies greater than 84% (Table 1). Our results corroborates previous findings on the diagnostic potential of metabolomics in heart failure [Cheng et al., 2015]and expands on them by employing a broader range of classification algorithms to achieve superior prediction accuracies [Marcinkiewicz-Siemion et al., 2020]. With further validation in prospective cohorts, these models could lead to the development of clinical tools leading to early diagnosis, paving the way for significantly enhance patient management and outcomes.

Machine learning models combined with explainability approaches offer significant potential for biomarker discovery, enabling the identification of molecular signatures associated with diseases. In the context of heart failure, the development of metabolomics signatures can reveal valuable biomarkers for diagnosis and prognosis. We illustrated this potential using a panel of metabolite measurements and identified a metabolic signature of HF samples through explainability analyses. We demonstrated that our XGBoost models were capable of reducing the phenotype signature to a concise list of metabolite markers, while the SVM model exhibited a higher degree of variability in global and local importance scores, resulting in a noisier output metabolite markers list. XGBoost models may thus be particularly useful for narrowing down noisy data to a sharp molecular signature of heart failure, with potential opportunities for exploring signature for other diseases.

Although the signature’s potential for prognosis in heart failure remains to be validated, it demonstrates our approach’s capability to gain insights into the biological mechanisms underlying this condition. Specifically, our workflow identified lignoceric acid (C24:0 FA), a very long-chain fatty acid (VLCFA), as a novel and significant discriminator. This result aligns with previous work showing that higher levels of VLCFA including lignoceric acid but also behenic acid have been shown to be associated with a lower risk of HF [Lemaitre et al., 2018] but also with lower risk of atrial fibrillation [Dinesen et al., 2018] or a decreased risk of coronary artery disease (Malik et al. [2015]). More recently, in a cohort of 4132 individuals from the 2003-2004, 2011-2012 National Health and Nutrition Examination Survey (NHANES), it has been shown that individuals with higher levels of VLCFA and particularly C22 and C24, showed a reduced risk of cardiovascular disease (CVD) or coronary heart disease (CHD) and even more mortality of all cause [Tao et al., 2024]. We then importantly replicated the differences of C24:0 FA levels between controls and heart failure patients in a replication cohort. To our knowledge, this is the first study to utilize metabolomics combined with explainable machine learning to discover a novel metabolite marker in heart failure, which could become a target for therapeutic interventions. Along with HDL cholesterol, glucose and C18:1 carnitine, lignoceric acid contributes to the metabolic signature of HFrEF.

Interaction analyses can deepen our understanding of the novel signature as a biological network. In the machine learning context, interactions refer to the combined effects of variables that improve the predictive power of models. In contrast, biological interactions involve the complex interplay between molecules within biological systems, reflecting underlying physiological processes. Our study identified an interacting network of metabolites responsible for heart failure by examining two-way variable interactions that collectively enhance the model’s predictions (Figure 4). This subset included the key metabolites presented above, as well as additional ones such as citrate and C22:6n3 fatty acid, highlighting the importance of considering non-linear relationships in metabolic networks [Mosconi et al., 2008]. Specifically, in the context of heart failure, the interactions among these metabolite markers provide a more accurate picture of the alterations occurring in the disease state, emerging from the contributions of several metabolites, each exhibiting distinct signatures across samples. The visual representation of our interaction network also offers insights into the complex biological mechanisms underlying the heart failure phenotype, and may also improve the identification of specific disease sub-phenotypes [Triposkiadis et al., 2019]. Adopting such integrative methods that emphasize molecular interactions should encourage further applications to uncover hidden relationships in other complex phenotypes.

The relatively small sample size of 124 samples and the fact all HF patients had a specific HF subtype limit the generalizability of our findings to all HF types, which is a limitation of our study. Future studies should determine whether our findings are specific to HFrEF or if they can be extended to HFmrEF and HFpEF. Moreover, future studies with larger sample sizes should test deep learning models that, while often deemed less explainable, can learn from large datasets more effectively [LeCun et al., 2015]. A more ethnically diverse dataset will also be necessary to validate the identified metabolite markers across different human populations. Another limitation is that our dataset consisted of 55 metabolites after applying all quality control steps, which may not fully capture the heterogeneity of heart failure. Having a few well-described biomarkers is however advantageous in clinical settings. Nevertheless, expanding the metabolite panel with untargeted metabolomics and incorporating other omics data (e.g., genetics, transcriptomics, proteomics) could provide a more comprehensive understanding of disease mechanisms. The metabolomics analysis, conducted solely on plasma samples at a single time point, provides a limited snapshot that does not capture pathway dynamics or establish causality. Finally, prospective validation is needed to demonstrate that the identified metabolic signature, as well as its effectiveness in assessing disease severity, can be reliably used for early diagnosis and patient management in real-world clinical settings.

In conclusion, we proposed an XAI workflow for biomarker discovery, combining metabolomics and machine learning. Our work serves as a proof of concept for the benefits of XAI in metabolomics, to elucidate the complex biological mechanisms underlying disease. The observed complex relationships between metabolites across patients reinforces the concept of precision medicine, highlighting the need for further exploration that could lead to significant advancements in heart failure diagnosis and treatment.

## Author contributions statement

CB (Conceptualization [lead], Formal analysis [lead], Investigation [lead], Methodology [lead], Software [lead], Validation [lead], Visualization [lead], Writing—original draft [lead]), P.M (Data curation [supporting], Methodology [supporting], Software [supporting], Writing—review & editing [supporting]), CD,LH,DB,JCD & JD (Data acquisition for validation [lead]), CDR (Data acquisition [lead], Conceptualization [supporting], Funding acquisition [lead], Supervision [supporting], Writing—review & editing [supporting]). MR (Data acquisition [lead], Conceptualization [supporting], Funding acquisition [lead], Supervision [supporting], Writing—review & editing [supporting]) and JGH (Conceptualization [lead], Funding acquisition [lead], Supervision [lead], Writing—review & editing [lead]). All authors revised and approved the final version of the manuscript.

## Acknowledgments

We thank members of the Hussin group for helpful discussions and advice throughout this project, specifically Raphael Poujol and Jean-Christophe Grenier. This work was completed thanks to computational resources provided by Calcul Quebec clusters Narval, Beluga, maintained by the Digital Research Alliance of Canada. This study was supported by funding from the Montreal Heart Institute Foundation, an IVADO PRF Grant to JGH and MR (PRF-2019-3378524797), a National Sciences and Engineering Research Council (NSERC) Discovery Grant to JGH (RGPIN-2022-04262), a Canadian Institutes of Health Research (CIHR) with contributions from the Institute of Nutrition, Metabolism and Diabetes to CDR (MA-9575), and an Agilent Technologies Research Grant to CDR (3893 & 4075). CB was supported by a PhD Scholarship from the NSERC-CREATE Metabolomics Advanced Training and International Exchange (MATRIX) program. MR is a Fonds de recherche du Québec en Santé (FRQS) Junior 2 research scholar (278281), JGH is FRQS Junior 2 research scholar (311067).

## Supplementary content

### Training reliable models for explainability analyses

As each fold might contain fold-specific explainability profiles, here the objective was to obtain a single model explainability to filter out any potential batch effects arising from fold sample composition. We hypothesized that the explainability derived from each fold is not different from the ones of models trained on the entire dataset, suggesting that the single model explainability would be as reliable as those of the individual folds, but more practical due to its singular nature. We first confirmed that the importance scores for each fold did not differ significantly from that of the models trained on the entire dataset. Indeed, the LIME weights derived from the 5-fold cross-validation models were compared with those from the model trained on the full dataset (referred to as *”full.model”*). These weights were positively correlated for both XGBoost(Pearson R = 0.68, p-value = 2.2E-16) and SVM (Pearson R = 0.79, p-value = 2.2E-16) models, indicating a high level of consistency in feature importance identified (Supplementary Figure S2). This illustrates that while some local explainability profiles vary depending on the fold composition, the feature importance in the full models is reliable, thus validating the approach of the *”full.model”* explainability presented in the subsequent analyses.

### Supplementary tables

**Table S1.** Dataset Overview.

| Dataset details | Before | After filtering and processing steps |
| --- | --- | --- |
| Metabolomics Protocols | Combination of targeted mass spectrometry-based methods and gas-chromatography with flame ionization detector (GC-FID) |  |
| Samples Origin | Plasma |  |
| Classes | Control / Heart failure with reduced ejection fraction (HFrEF) |  |
| Sample size | 132 (Control = 72, HFrEF = 60) | 124 (Control = 71, HFrEF = 53) |
| Measured Metabolites | 71 | 55 |
| Study | Asselin et al [Asselin et al., 2014] - Ruiz et al [Ruiz et al., 2017] |  |

Table summarizing the metabolomics dataset used before and after quality control steps indicated in each column. Metabolomics Protocols: describes the technical methods used to obtain the metabolomics data. Samples origin: indicates the biological source of the samples. Classes: shows the two groups compared in the study: control samples and patients with heart failure. Sample size: total number of samples with a specification of the number of control samples and number HFrEF patient samples. Measured metabolites: number of metabolites. Study: cites the studies in which the data was originally reported. Abbreviations : HFrEF = heart failure with reduced ejection fraction.

**Table S2.** Dataset overview for the replication cohort.

|  | **Controls** | **HF** | **HFrEF** |
| --- | --- | --- | --- |
| **n** | 71 | 31 | 14 |
| **Age.Years (mean (SD))** | 66.04 (8.59) | 70.10 (11.82) | 67.86 (13.23) |
| **Sex = 1 (%)\*** | 42 (59.2) | 14 (45.2) | 9 (64.3) |
| **NT-proBNP (mean (SD))** | NaN (NA) | 2053.03 (1572.08) | 2712.07 (1596.78) |
| **eGFR (mean (SD))** | NaN (NA) | 37.20 (10.41) | 37.97 (11.85) |
| **LVEF (mean (SD))** | NaN (NA) | 44.13 (14.91) | 30.07 (7.92) |

Table summarizing the demographic and clinical characteristics of the study samples in the replication cohort, stratified by control group (Controls), heart failure group (HF), and heart failure with reduced ejection fraction group (HFrEF). The number of participants (n) in each group is listed at the top. The control group data serve as a baseline for comparison, although NT-proBNP, eGFR, and LVEF values are not reported (NaN) for this group. The values presented in the table represent means, with standard deviations (SD) shown in parentheses. \*The number and percentage of male participants in each group. Abbreviations - eGFR : estimated Glomerular Filtration Rate, FA : Fatty Acid, LVEF : Left Ventricle Ejection Fraction, NT-proBNP : N-terminal pro B-type Natriuretic Peptide.

**Table S3.** Overview of model performances, for the 5 cross-validation schemes, across 3 different seeds.

| Performance Metric Name | Logit | SVM | XGBoost |
| --- | --- | --- | --- |
| **Seed 124** |  |  |  |
| Specificity | 83.2 - 8.44 | 90.2 - 7.98 | 83.2 - 3.90 |
| Sensitivity | 77.6 - 12.22 | 79.8 - 9.31 | 84.6 - 10.57 |
| Balanced Accuracy | 80.2 - 3.27 | 84.8 - 8.01 | 84.0 - 6.71 |
| Accuracy | 80.6 - 2.88 | 85.8 - 8.26 | 84.0 - 7.38 |
| **Seed 125** |  |  |  |
| Specificity | 89.0 - 9.14 | 91.8 - 5.63 | 74.0 - 13.21 |
| Sensitivity | 83.4 - 11.44 | 76.0 - 13.21 | 90.2 - 6.26 |
| Balanced Accuracy | 86.0 - 5.15 | 83.8 - 6.18 | 81.8 - 7.66 |
| Accuracy | 86.4 - 5.22 | 85.0 - 5.92 | 83.2 - 7.46 |
| **Seed 126** |  |  |  |
| Specificity | 87.2 - 11.24 | 91.6 - 3.13 | 88.8 - 3.83 |
| Sensitivity | 83.0 - 8.43 | 79.6 - 7.09 | 85.8 - 12.56 |
| Balanced Accuracy | 85.2 - 7.85 | 85.4 - 4.83 | 87.2 - 8.11 |
| Accuracy | 85.6 - 8.29 | 86.4 - 4.56 | 87.2 - 8.67 |
| **Mean - SD across Seeds** |  |  |  |
| Specificity | 86.47 - 9.61 | 91.20 - 5.58 | 82.00 - 6.98 |
| Sensitivity | 81.33 - 10.70 | 78.47 - 9.87 | 86.87 - 9.80 |
| Balanced Accuracy | 83.80 - 5.42 | 84.67 - 6.34 | 84.33 - 7.49 |
| Accuracy | 84.20 - 5.46 | 85.73 - 6.25 | 84.80 - 7.84 |

**Table S4.** Variable Selection Logit models.

| Selected | Fold 1 |  | Fold 2 |  | Fold 3 |  | Fold 4 |  | Fold 5 |  |
| --- | --- | --- | --- | --- | --- | --- | --- | --- | --- | --- |
|  | Name | Score | Name | Score | Name | Score | Name | Score | Name | Score |
| **Seed 124** |  |  |  |  |  |  |  |  |  |  |
| Metabolite 1 | HDL-cholesterol | 0.67 | HDL-cholesterol | 0.58 | HDL-cholesterol | 0.60 | C24:0_FA | 0.75 | C24:0_FA | 0.75 |
| Metabolite 2 | C24:0_FA | 0.53 | C24:0_FA | 0.49 | C24:0_FA | 0.51 | HDL-cholesterol | 0.64 | HDL-cholesterol | 0.74 |
| Metabolite 3 | C18:1_car | 0.42 | total-cholesterol | 0.42 | C22:6n3_FA | 0.41 | C22:6n3_FA | 0.48 | C22:6n3_FA | 0.43 |
| Metabolite 4 | C22:6n3_FA | 0.39 | C18:1n7_FA | 0.41 | C20:5n3_FA | 0.36 | C20:5n3_FA | 0.38 | C22:0_FA | 0.43 |
| Metabolite 5 | LDL-cholesterol | 0.38 | LDL-cholesterol | 0.38 | C18:1n7_FA | 0.35 | LDL-cholesterol | 0.38 | glucose | 0.42 |
| Metabolite 6 | total-cholesterol | 0.34 | C22:6n3_FA | 0.37 | total-cholesterol | 0.32 | C22:0_FA | 0.34 | C20:5n3_FA | 0.35 |
| Metabolite 7 | C18:1n9_FA | 0.34 | citrate | 0.35 | C22:0_FA | 0.30 | total-cholesterol | 0.33 | total-cholesterol | 0.34 |
| Metabolite 8 | C20:5n3_FA | 0.33 | C20:5n3_FA | 0.34 | C18:1_car | 0.27 | C18:1_car | 0.30 | citrate | 0.30 |
| Metabolite 9 | glucose | 0.31 | C18:1_car | 0.33 | citrate | 0.27 | C2_car | 0.28 | LDL-cholesterol | 0.29 |
| Metabolite 10 | C22:0_FA | 0.30 | glucose | 0.29 | LDL-cholesterol | 0.25 | citrate | 0.24 | C18:1_car | 0.28 |
| **Seed 125** |  |  |  |  |  |  |  |  |  |  |
| Metabolite 1 | HDL-cholesterol | 0.56 | HDL-cholesterol | 0.67 | HDL-cholesterol | 0.65 | C24:0_FA | 0.72 | HDL-cholesterol | 0.62 |
| Metabolite 2 | C24:0_FA | 0.53 | C24:0_FA | 0.61 | C24:0_FA | 0.57 | HDL-cholesterol | 0.72 | C24:0_FA | 0.55 |
| Metabolite 3 | C22:6n3_FA | 0.46 | C22:6n3_FA | 0.38 | LDL-cholesterol | 0.38 | C22:6n3_FA | 0.50 | C22:6n3_FA | 0.38 |
| Metabolite 4 | total-cholesterol | 0.36 | total-cholesterol | 0.34 | C20:5n3_FA | 0.38 | C22:0_FA | 0.49 | LDL-cholesterol | 0.36 |
| Metabolite 5 | C20:5n3_FA | 0.34 | citrate | 0.33 | C22:6n3_FA | 0.38 | C18:1_car | 0.42 | C18:1n7_FA | 0.35 |
| Metabolite 6 | C18:1_car | 0.33 | C22:0_FA | 0.33 | total-cholesterol | 0.36 | C20:5n3_FA | 0.40 | C18:1_car | 0.32 |
| Metabolite 7 | C22:0_FA | 0.30 | C20:5n3_FA | 0.33 | citrate | 0.32 | total-cholesterol | 0.38 | glucose | 0.31 |
| Metabolite 8 | LDL-cholesterol | 0.30 | LDL-cholesterol | 0.31 | glucose | 0.31 | LDL-cholesterol | 0.31 | total-cholesterol | 0.31 |
| Metabolite 9 | glucose | 0.28 | C18:1n7_FA | 0.30 | C22:0_FA | 0.30 | glucose | 0.30 | C20:5n3_FA | 0.31 |
| Metabolite 10 | citrate | 0.24 | C18:1_car | 0.29 | C18:1n7_FA | 0.29 | C18:1n7_FA | 0.28 | citrate | 0.30 |
| **Seed 126** |  |  |  |  |  |  |  |  |  |  |
| Metabolite 1 | HDL-cholesterol | 0.61 | HDL-cholesterol | 0.62 | HDL-cholesterol | 0.81 | C24:0_FA | 0.65 | HDL-cholesterol | 0.56 |
| Metabolite 2 | C24:0_FA | 0.51 | C24:0_FA | 0.60 | C24:0_FA | 0.67 | HDL-cholesterol | 0.63 | C24:0_FA | 0.56 |
| Metabolite 3 | LDL-cholesterol | 0.37 | C22:6n3_FA | 0.47 | C22:6n3_FA | 0.51 | C22:6n3_FA | 0.45 | total-cholesterol | 0.32 |
| Metabolite 4 | C22:6n3_FA | 0.37 | total-cholesterol | 0.38 | C20:5n3_FA | 0.44 | C22:0_FA | 0.43 | C18:1n7_FA | 0.32 |
| Metabolite 5 | citrate | 0.34 | C20:5n3_FA | 0.38 | C22:0_FA | 0.36 | C20:5n3_FA | 0.40 | C22:6n3_FA | 0.31 |
| Metabolite 6 | total-cholesterol | 0.33 | C18:1n7_FA | 0.34 | total-cholesterol | 0.35 | total-cholesterol | 0.37 | glucose | 0.30 |
| Metabolite 7 | C18:1_car | 0.33 | C18:1_car | 0.33 | LDL-cholesterol | 0.34 | C18:1_car | 0.33 | LDL-cholesterol | 0.30 |
| Metabolite 8 | glucose | 0.32 | LDL-cholesterol | 0.33 | C18:1_car | 0.33 | LDL-cholesterol | 0.32 | citrate | 0.29 |
| Metabolite 9 | C20:5n3_FA | 0.30 | C22:0_FA | 0.31 | glucose | 0.30 | citrate | 0.28 | C22:0_FA | 0.29 |
| Metabolite 10 | C22:0_FA | 0.27 | glucose | 0.30 | C2_car | 0.29 | C18:1n7_FA | 0.26 | C18:1_car | 0.26 |

This table presents the selected metabolites based on their corresponding F-scores across five folds. Each fold represents a cross-validation iteration, listing the metabolites chosen for fitting the models. The rows indicate the seed values used for each set of logit models.

**Table S5.** Grid Search results for SVM models.

| cost | gamma | seed_value | fold |
| --- | --- | --- | --- |
| 8.2 | 0.001 | 124 | 1 |
| 8.4 | 0.0005 | 124 | 2 |
| 5.0 | 0.005 | 124 | 3 |
| 8.0 | 0.0005 | 124 | 4 |
| 6.6 | 0.001 | 124 | 5 |
| 5.6 | 0.001 | 125 | 1 |
| 9.8 | 0.001 | 125 | 2 |
| 5.0 | 0.0005 | 125 | 3 |
| 6.8 | 0.001 | 125 | 4 |
| 5.0 | 0.0005 | 125 | 5 |
| 6.0 | 0.001 | 126 | 1 |
| 9.8 | 0.001 | 126 | 2 |
| 5.0 | 0.0005 | 126 | 3 |
| 6.8 | 0.005 | 126 | 4 |
| 5.6 | 0.0005 | 126 | 5 |

This table presents the combinations of hyperparameters used in the SVM models. Each row represents a unique fold with a specific set of hyperparameters, including "cost" and "gamma". The possible values for hyperparameter tuning are: "cost" can take values from the sequence generated by 10 raised to the power of -1 to 2, resulting in values 0.1, 1, 10, 100 "gamma" from a starting value of 0.5 to 4, in increments of 0.10. A total of 15 rows, 5 folds for three different seed values (124, 125, 126). Abbreviations : SVM = Support-Vector Machine.

**Table S6.** Grid Search results for XGBoost models.

| nround | max_depth | eta | gamma | colsample | min_child | sub_sample | seed_value | fold |
| --- | --- | --- | --- | --- | --- | --- | --- | --- |
| 1000 | 5 | 0.4 | 0 | 1 | 0.5 | 1 | 124 | 1 |
| 1000 | 1 | 0.6 | 0 | 1 | 0.5 | 1 | 124 | 2 |
| 1000 | 1 | 0.8 | 2 | 1 | 0 | 1 | 124 | 3 |
| 1000 | 1 | 0.4 | 0 | 1 | 0 | 1 | 124 | 4 |
| 1000 | 5 | 0.6 | 0 | 1 | 1 | 1 | 124 | 5 |
| 1000 | 1 | 1 | 0 | 1 | 0.5 | 1 | 125 | 1 |
| 1000 | 5 | 0.2 | 2 | 1 | 0 | 1 | 125 | 2 |
| 1000 | 5 | 0.6 | 0 | 1 | 1 | 1 | 125 | 3 |
| 1000 | 5 | 0.4 | 1 | 1 | 0 | 1 | 125 | 4 |
| 1000 | 5 | 0.6 | 2 | 1 | 2 | 1 | 125 | 5 |
| 1000 | 5 | 0.8 | 2 | 1 | 2 | 1 | 126 | 1 |
| 1000 | 5 | 1 | 1 | 1 | 1 | 1 | 126 | 2 |
| 1000 | 5 | 0.6 | 1 | 1 | 1 | 1 | 126 | 3 |
| 1000 | 1 | 0.2 | 0 | 1 | 0 | 1 | 126 | 4 |
| 1000 | 1 | 1 | 1 | 1 | 0 | 1 | 126 | 5 |

Table presents the combinations of hyperparameters used in the XGBoost models. Each column represents a unique fold with a specific set of hyperparameters, including "nround", "max\_depth", "eta", "gamma", "colsample", "min\_child", "sub\_sample". The possible values for hyperparameter tuning are: "nrounds" is fixed at 1000, "eta" can take any value from the set 0, 0.2, 0.4, 0.6, 0.8, 1.0, "max\_depth" from 1, 5, 10, 15, 20, "gamma" from 0, 1, 2, 5, 10, "colsample\_bytree" is fixed at 1, "min\_child\_weight" from 0, 0.5, 1, 2, 5, and "subsample" is fixed at 1. A total of 15 rows, 5 folds for three different seed values (124, 125, 126). Abbreviations : XGBoost = eXtreme Gradient Boosting.

### Supplementary figures

**Fig. S1.**
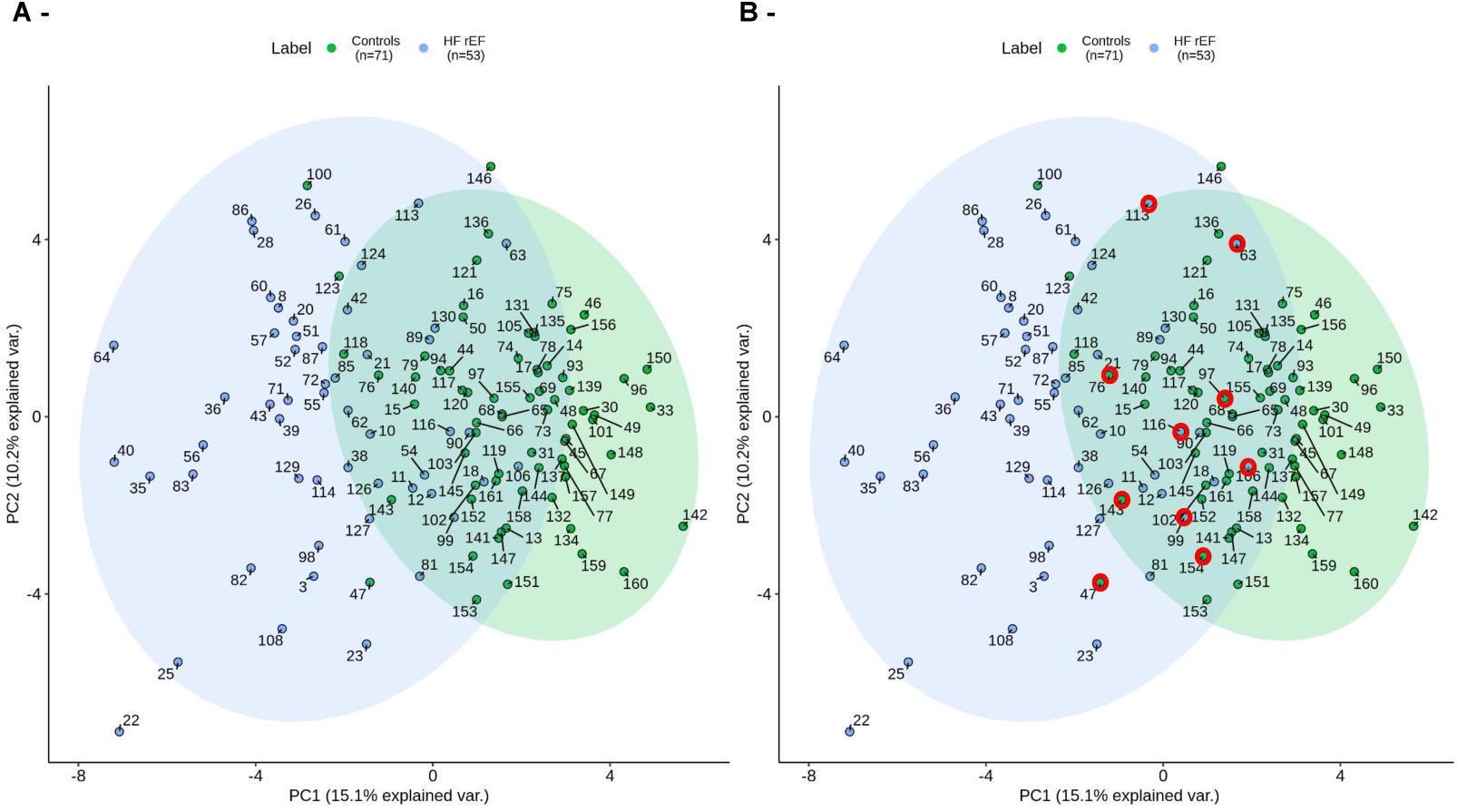
Principal Component Analysis (PCA) of the 124 samples and the 55 variables. Principal component analysis (PCA) of 124 samples (71 controls, 53 HF patients) using 55 variables adjusted for age and sex. Controls are identified in green, and HF patients in blue. A: Quality control using. B: PCA highlighting XGBoost misclassified samples across three seeds and 5-CV. Abbreviations: HFrEF = heart failure with reduced ejection fraction, var = variance.

**Fig. S2.**
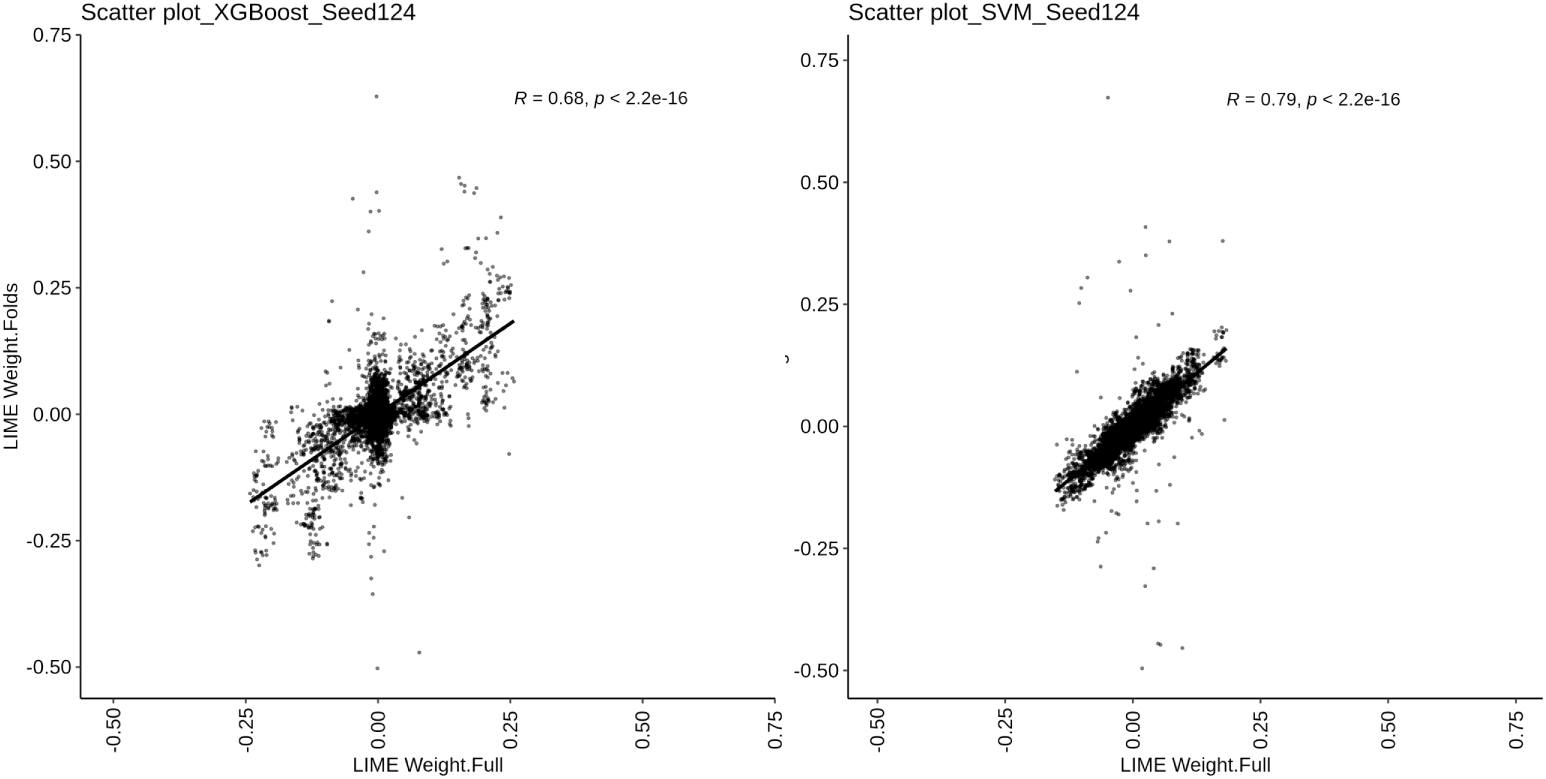
LIME weights comparison between 5-cross validation models and full models. Scatter plots of LIME weights. (Left) XGBoost model with seed 124. (Right) SVM model with seed 124.

**Fig. S3.**
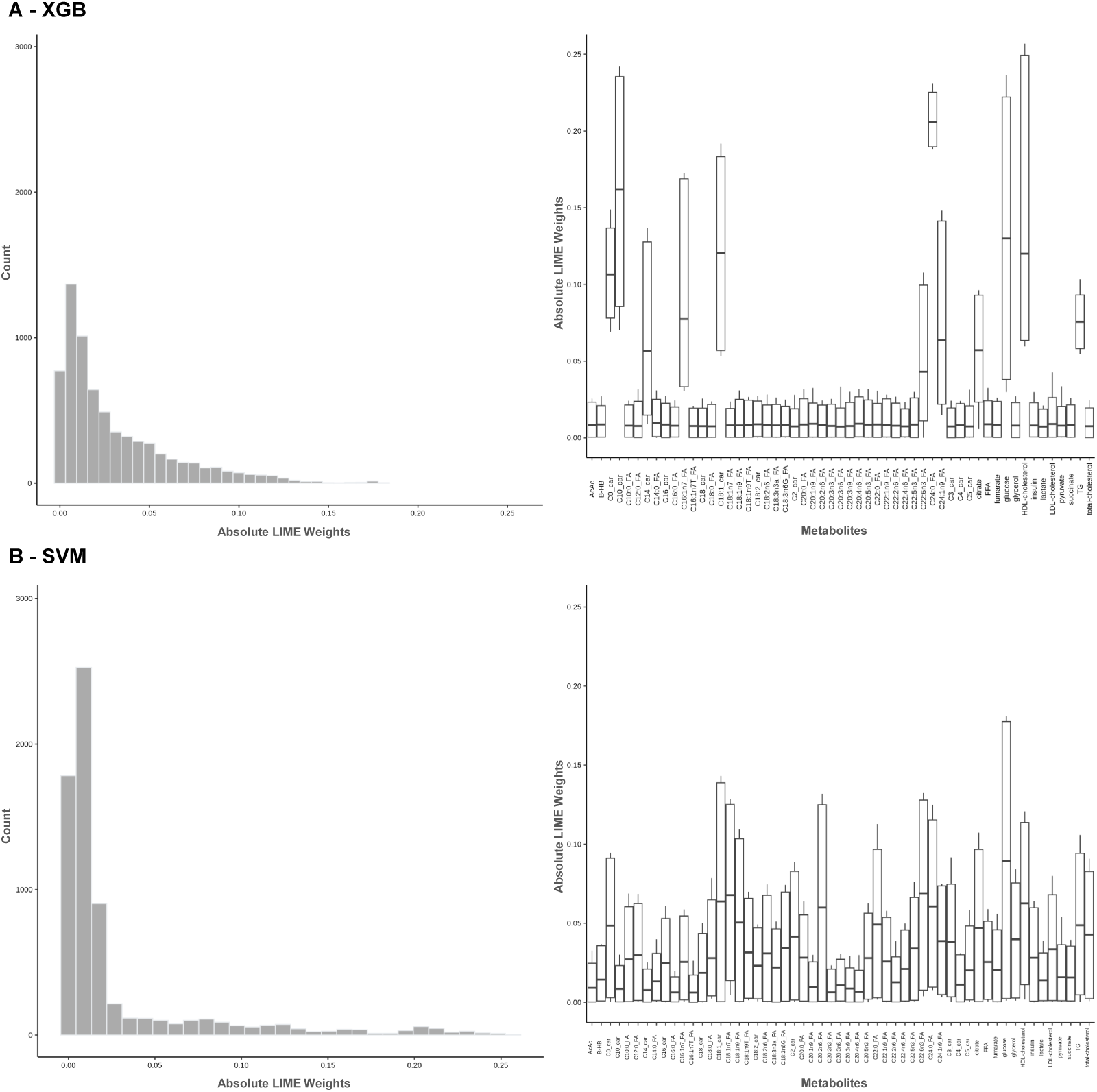
LIME weights distribution comparison between SVM and XGB. Total and metabolite distribution of absolute LIME weights for XGBoost (A) and SVM (B) models. Left panels - Histogram of absolute LIME weights for XGBoost, indicating the frequency of weights assigned to metabolites. Right panels - Boxplots showing the distribution of LIME weights per metabolite. Abbreviations - SVM = Support-Vector Machine, XGBoost = eXtreme Gradient Boosting, FA : Fatty Acid, car : carnitine.

**Fig. S4.**
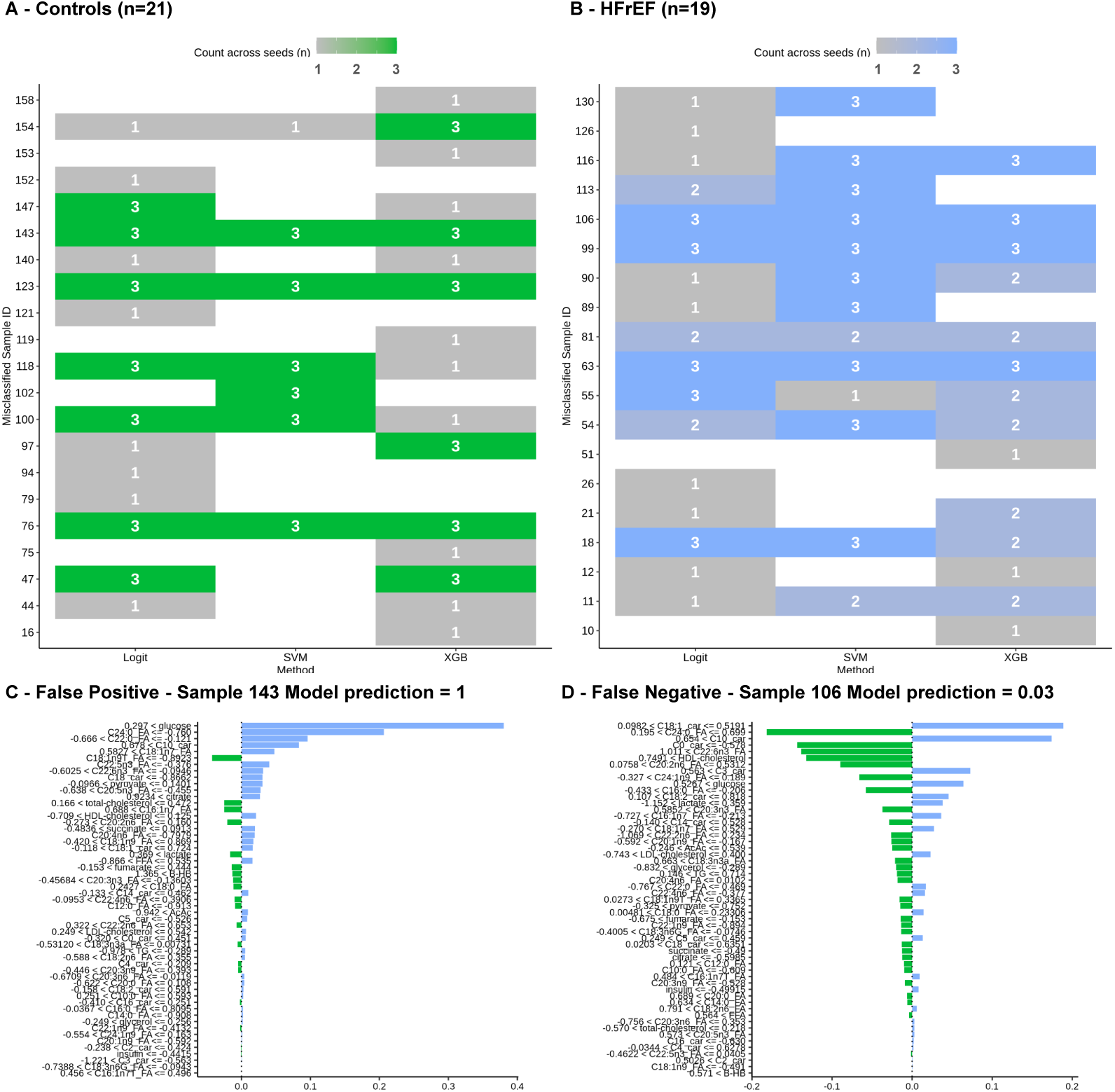
List of misclassified samples across various models. List of misclassified samples across Logit, SVM, XGBoost models for each seed (124,125,126). The count is representing the total occurrence of the wrong classification. The maximum possible value is 3 as each sample is tested once per seed. A: control samples. B: HFrEF samples. C: The false positive with the largest prediction error. D: The false negative with the largest prediction error.

**Fig. S5.**
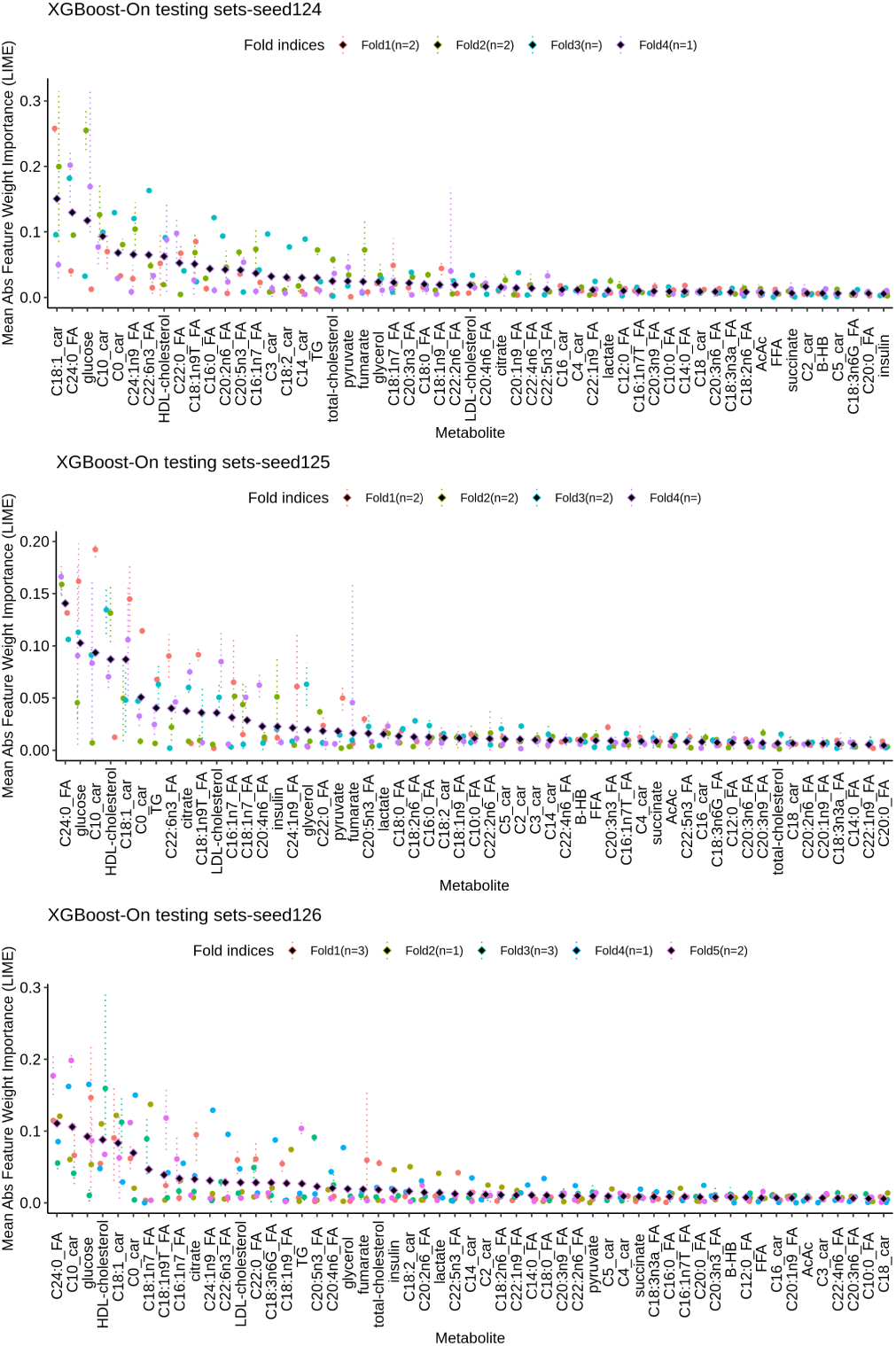
Wrongly classified samples for XGBoost model, important variables. Dot plots displaying the mean absolute LIME weights for each metabolite across the folds of the XGBoost models. Each dot is color-coded according to the fold, with dotted lines indicating the minimum and maximum absolute LIME weights for the misclassified samples within a specific fold. The diamond shape represents the mean LIME weight for each metabolite across the different folds. Abbreviations used: XGBoost = eXtreme Gradient Boosting, FA = Fatty Acid, car = carnitine.

**Fig. S6.**
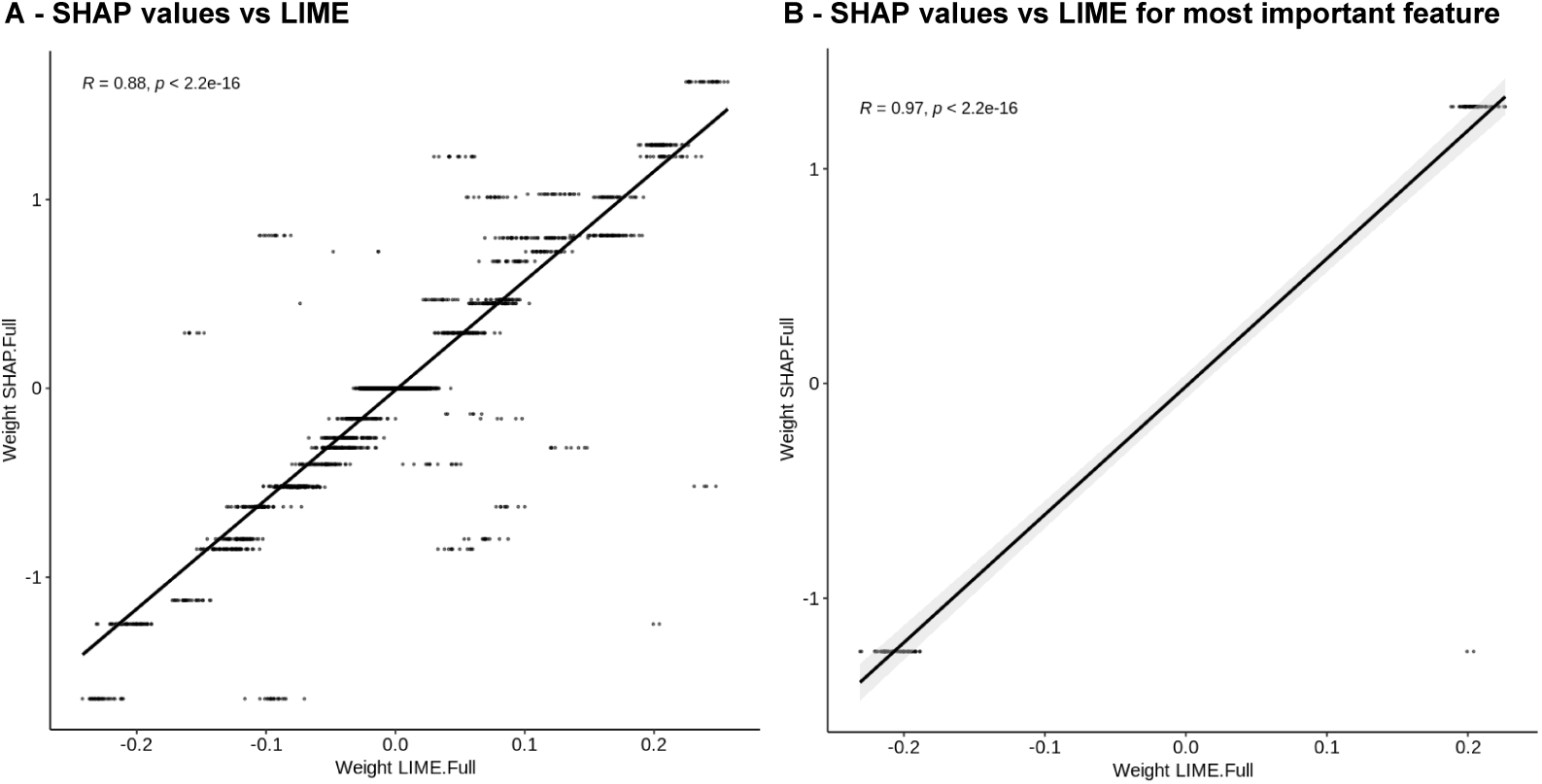
Local feature weights distribution comparison between SHAP values and LIME for the full model XGBoost. Scatter plots of across all samples and metabolites (A) and specifically for the most discriminant metabolite, namely C24:0 FA (B). Abbreviations - XGBoost = eXtreme Gradient Boosting, FA : Fatty Acid.

